# A Mesenchymal Tumor Cell State Confers Increased Dependency on the BCL-X_L_ Anti-apoptotic Protein in Kidney Cancer

**DOI:** 10.1101/2022.01.29.478337

**Authors:** Treg Grubb, Smruthi Maganti, John Michael Krill-Burger, Cameron Fraser, Laura Stransky, Tomas Radivoyevitch, Kristopher A. Sarosiek, Francisca Vazquez, William G. Kaelin, Abhishek A. Chakraborty

**Author notes:** These authors contributed equally (Treg Grubb, Smruthi Maganti). Correspondence should be addressed to W.G.K and A.A.C.

## Abstract

Genome-wide genetic screens have identified cellular dependencies in many cancers. Using the Broad Institute’s Achilles shRNA screening dataset, we mined for targetable dependencies by cell lineage. Our studies identified a strong dependency on *BCL2L1*, which encodes the BCL-X_L_ anti-apoptotic protein, in a subset of kidney cancer cells. Genetic and pharmacological inactivation of BCL-X_L_, but not the related anti-apoptotic proteins BCL-2, led to fitness defects in renal cancer cells, and also sensitized them to chemotherapeutics. Neither BCL-X_L_ levels (absolute or normalized to BCL-2) nor the status of the *VHL* gene, which is frequently mutated in kidney cancer, predicted BCL-X_L_ dependence. Transcriptional profiling, however, identified a ‘BCL-X_L_ dependency’ mRNA signature, which included elevated mesenchymal gene expression in BCL-X_L_ dependent cells. Promoting mesenchymal transition increased BCL-X_L_ dependence; whereas, conversion to a more differentiated state overcame BCL-X_L_ dependence in kidney cancer cells. The ‘BCL-X_L_ dependency’ mRNA signature was observed in almost a third of human clear cell Renal Cell Carcinomas (ccRCCs), which were also associated with worse clinical outcomes. Finally, an orally bioavailable BCL-X_L_ inhibitor, A-1331852, showed anti-tumor efficacy *in vivo*. Altogether, our studies uncovered an unexpected link between cancer cell state and dependence on the anti-apoptotic BCL-X_L_ protein and justify further testing on BCL-X_L_ blockade as a potential way to target a clinically aggressive subset of human kidney cancers.

**One Sentence Summary:** Cell state, but not pVHL and/or HIF status, defines the dependency of kidney cancer cells on the BCL-X_L_ anti-apoptotic protein.

## INTRODUCTION

Renal Cell Carcinoma (RCC) ranks among the top ten forms of cancer in humans, with current estimates projecting ~79,000 new diagnoses and ~14,000 deaths this year in USA (*1*). Early stage RCC, which is observed in nearly 60% of patients, can be effectively managed with either surgery or surveillance; however, nearly 4 in 10 patients present with advanced or metastatic RCC, and experience much poorer outcomes. Moreover, almost a third of the patients with higher-risk localized disease during initial diagnosis subsequently develop metastatic disease. Therapeutic options to manage advanced disease presently include combining a checkpoint inhibitor (CKI) either with another CKI or with a tyrosine kinase inhibitor (TKI), and the most recent trials have shown that between 30-60% patients exhibit measureable responses with these therapeutic combinations (*2*–*4*). However, there is limited recourse for the remaining patients. Finally, most patients eventually develop resistance to these treatments and subsequent treatment options are largely ineffective.

The clear cell Renal Cell Carcinomas (ccRCCs) represent ~70-75% of RCCs (*5*). Inactivation of the von Hippel-Lindau tumor suppressor protein (pVHL) is a nearly universal hallmark of ccRCC (*5*), and mechanistic studies defining the consequences of pVHL loss have informed the pre-clinical discovery of actionable dependencies in ccRCC (*6*–*8*). pVHL functions as the substrate recognition subunit in an E3 ligase complex that is perhaps best known to mediate oxygen-dependent destruction of the α-subunit of the Hypoxia Inducible Factor (HIFα) (*9, 10*). Consequently, HIFα accumulates in pVHL-deficient ccRCCs, independent of oxygen availability. There are two transactivation-competent isoforms of HIFα (HIF1α and HIF2α) and chronic activation of HIF2α drives kidney cancer pathogenesis (*11, 12*). Therapeutic strategies to target HIF2α directly have been developed in the last decade (*13*, *14*). These approaches have shown promise in both pre-clinical (*15*, *16*) and clinical settings (*17*, *18*). But not all patients respond, and not all responses are durable. Therefore, new treatments, especially to target tumors that are refractory to HIF2α inhibitors (and other therapies), are urgently needed.

Recent largescale screening efforts, such as the Broad Institute’s Achilles (*19*) and DepMap projects (*20*), and Novartis’ DRIVE project (*21*), have provided investigators with publically accessible resources that enable the identification of novel oncogenic dependencies. These screening strategies rely on either genetic (e.g. shRNA and CRISPR/Cas9) or pharmacological tools. These different tools can produce different phenotypes, even when directed at the same target. For example, CRISPR/Cas9-based screens routinely yield phenotypes associated with near complete elimination (knockout) of the gene product. In contrast, shRNA screens often score phenotypes associated with partial mRNA reduction (knockdown), and perhaps better model the incomplete blockade of target proteins that is typically achieved with drugs. Importantly, though, the candidates identified through shRNA-based approaches need to be rigorously vetted because of the notorious “off-target” cytotoxicity associated with shRNA use (*22*).

Here, we exploited the availability of multiple genetic screening databases to identify novel targetable dependencies in kidney cancer. We reasoned that the genetic lesions acquired during renal oncogenesis and/or the master regulators of lineage specification could confer targetable dependencies in these cancer cells. Therefore, a comparison of genetic dependencies in the kidney lineage cells could identify targetable vulnerabilities that are preferentially enriched in RCC (versus other lineages).

## Results

### BCL-X_L_ is a lineage-specific dependency in kidney cancer

To identify dependencies that are selectively enriched in kidney cancer cells (versus all other lineages), we used the Broad Institute’s (shRNA-based) Achilles dataset. This dataset measures the enrichment or depletion of shRNAs in a large representative sample set of cancer cells representing various cancer types and lineages. The cell lines in this dataset were lentivirally transduced in a pooled format with a genome-wide shRNA library and cultured over 6-8 weeks to track changes in the abundance of shRNAs by next-gen sequencing (*19*).

Given the concern of losing critical dependencies because they failed to score in all of the cell lines due to inherent genetic heterogeneity (i.e. false negatives), we sorted dependencies that showed strong selectivity in at least a subset of kidney lineage cells [Likelihood Ratio Test (LRT)>100]. We then excluded “common essential genes”, which typically have housekeeping roles, and confirmed that the scored dependency was observed in a minimum of three independent cell lines within the kidney lineage (Table S1), thereby scoring ~twenty potential dependencies. We then ranked the dependencies by LRT and interrogated the top ten “hits” from this analysis (Fig. 1A). We noted that the top two candidates were *HNF1B* and *PAX8*, both of which have an established role in kidney lineage specification (*23, 24*). Most of the other candidates on the list included ribosomal subunits, which are not inherently druggable; however, *BCL2L1*, scored as the top gene that encoded a druggable gene product. The impact of *BCL2L1* loss by shRNA on kidney cancer cells was confirmed using the more recent CRISPR/Cas9 dependency maps (*20*). We observed a great degree of similarity in *BCL2L1* dependence between the two datasets and confirmed that a subset of kidney cancer cells (e.g. CAKI-2, TUHR4TKB, and TUHR10TKB) showed very strong dependence on *BCL2L1* in both datasets (Fig. 1B).

**FIGURE 1.**
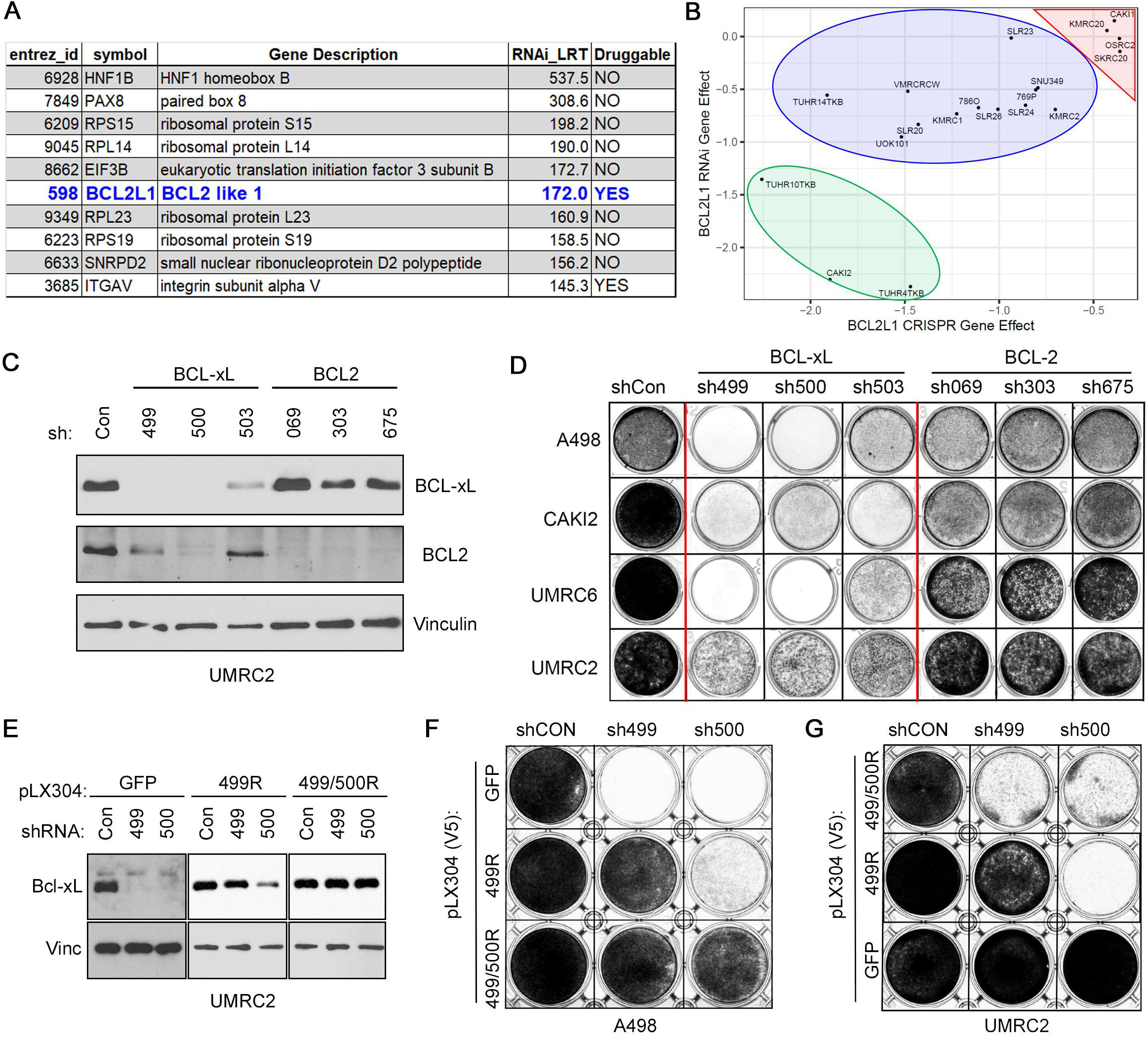
BCL-X_L_ is a Strong Dependency in a Subset of Kidney-lineage Cancer Cells. (**A** and **B**) List of dependencies enriched for selectivity in a subset of kidney lineage cells, as indicated by Likelihood Ratio Test (LRT) scores > 100 (**A**) and comparison of BCL-X_L_ dependency in the RNAi versus CRISPR/Cas9 dependency maps, in the indicated kidney cancer cell lines (**B**). Negative scores indicate a dependency with scores < −1 (strong, green oval), −0.5 to −1.0 (intermediate, blue oval), and >-0.5 (neutral/resistant, red triangle), annotated based on the CRISPR/Cas9 dataset. (**C** and **D**) Immunoblots (**C**) and crystal violet staining (**D**) of the indicated cell lines that were lentivirally transduced to express shRNAs targeting the indicated BCL-2-family genes or non-targeting controls (Con). (**E** to **G**) Immunoblots (**E**) and crystal violet staining (**F** and **G**) of the indicated cell lines that were lentivirally transduced to express the indicated shRNA-resistant versions of BCL-X_L_ or GFP, as a control, followed by lentiviral expression of the indicated shRNAs. (**C**) and (**E**) were done 3 days post infection with the shRNA-expressing lentiviral particles. (**D**), (**F**), and (**G**) were done 3 days post selection for A-498, CAKI-2, and UMRC-6 cells, and 7-10 days post selection for UMRC-2 cells.

The *BCL2L1* gene encodes the anti-apoptotic BCL-X_L_ protein, a member of the broader BCL-2-family of proteins, which includes BCL-2, BCL-w, MCL-1, and BFL-1/A1. To confirm our initial observations, we lentivirally transduced representative ccRCC cell lines (A-498, CAKI-2, UMRC-2, and UMRC-6) with three unique shRNAs that targeted either *BCL2L1* (499, 500, and 503), *BCL2* (069, 303, and 675), or non-targeting controls (shCon). After selection in puromycin, we stained the surviving cellular population with crystal violet. Knockdown of *BCL2L1* caused evident cytotoxicity in A-498, CAKI-2, and UMRC-6 within 3-5 days post selection and in UMRC-2 cells ~10-12 days post selection (Fig. 1, C and D). In contrast, *BCL2* shRNAs did not cause such profound cytotoxicity in the kidney cells, despite robustly knocking down BCL2 (Fig. 1, C and D).

Two BCL-X_L_ targeting shRNAs (499 and 500) also partially downregulated BCL-2. To formally establish that the cytotoxic effects of these *BCL2L1* shRNAs were “on-target”, we first engineered a BCL-X_L_ cDNA that harbored silent mutations in the sh499 recognition sequence. Expression of this construct (499R) rescued the cytotoxicity associated with sh499, but not sh500, in two different cell lines (Fig. 1, E to G). We then introduced additional silent mutations into the 499R backbone to also make this cDNA resistant to sh500 (499/500R). Expression of this BCL-X_L_ cDNA rescued the cytotoxicity associated with both sh499 and sh500 in both the renal cell lines (Fig. 1, E to G). Together, these genetic studies confirmed that ccRCCs are dependent on the BCL-X_L_ anti-apoptotic protein, but less so on the closely related BCL-2 protein; and indicated that the inherent sensitivity of ccRCCs might differ between cell lines, with cells like UMRC-2 presenting with delayed cytotoxicity in response to BCL-X_L_ loss.

### Pharmacological BCL-X_L_ blockade mimics results of genetic studies

The BCL-2-family anti-apoptotic proteins have been targets of clinical interest for many years and, as such, have been subjected to intensive “hit-to-lead” campaigns to identify highly specific pharmacological inhibitors. These efforts have identified the BCL-X_L_/BCL-2-dual inhibitors ABT-737 (*25*) and ABT-263 (Navitoclax) (*26*), the BCL-2-specific inhibitor ABT-199 (Venetoclax) (*27*), and more recently the BCL-X_L_-specific inhibitor A-1331852 (*28*).

To validate the findings from our genetic studies, we characterized a panel of ten ccRCC lines for sensitivity to the BCL-2-family inhibitors. We observed that cell lines that were exquisitely sensitive to BCL-X_L_ loss in genetic studies (e.g. CAKI-2 and TUHR4TKB) were also highly sensitive to three days of treatment with either ABT-263 (fig. S1) or the BCL-X_L_-specific inhibitor A-1331852, with cellular IC_50_ values in the low nM range (Fig. 2A and fig. S2A). In contrast, cell lines like UMRC-2, which showed delayed response to genetic BCL-X_L_ loss, were likewise less responsive to three days of treatment with pharmacological agents that target BCL-X_L_ (Fig. 2A and figs. S1 and S2A). These analyses also identified the A-498 and SLR23 cells as additional examples of A-1331852 sensitive ccRCCs. The BCL-X_L_ dependency in these cells was specific; because, in contrast to the BCL-2-dependent B-cell lymphoma line SU-DHL-6, the ccRCC lines were virtually resistant to the BCL-2 inhibitor ABT-199 (Fig. 2B and fig. S2B). Altogether, nearly 40% of the ccRCC lines were sensitive (IC_50_ in the nM range) to A-1331852. Importantly, sensitivity in only a subset of cell lines is a common occurrence, perhaps due to underlying genetic heterogeneity among cell lines. For instance, the HIF2α inhibitor, which has recently received FDA approval for use against ccRCCs, efficiently targets only a subset of ~20-30% ccRCCs.

**FIGURE 2.**
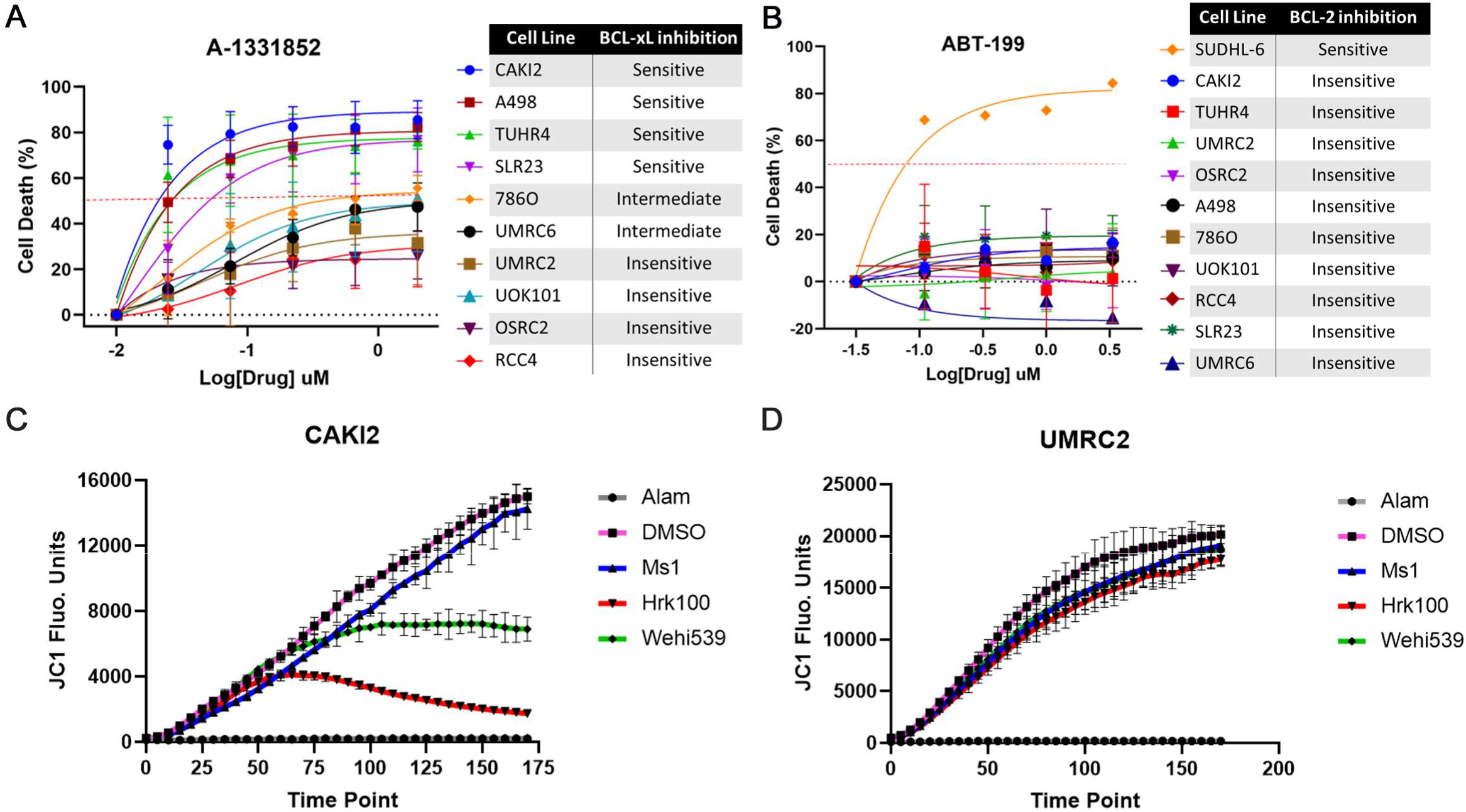
BCL-X_L_ Inhibition Promotes Apoptotic Cell Death in a Subset of ccRCCs, (**A** and **B**) Percent cell death, relative to the DMSO-treated control cells, determined using the XTT assay in the indicated ccRCC cell lines that were treated with the indicated concentrations of the BCL-X_L_ inhibitor A-1331852 (A) or the BCL-2 inhibitor ABT-199 (B) for 3 days. (**C** and **D**) JC1 fluorescence measurement at the indicated time-points in CAKI-2 (**C**) and UMRC-2 cells (**D**) that were exposed to Alamethicin (Alam) (positive control), DMSO (negative control), sensitizer BH3 peptides (MS1 targeting MCL-1 and Hrk targeting BCL-X_L_), or the small molecule BCL-X_L_ inhibitor Wehi539.

We confirmed that the cytotoxicity triggered by A-1331852 treatment occurred because of increased apoptosis. Upon A-1331852 treatment, Annexin V (AnnV) versus Propidium Iodide staining showed a significant increase in apoptotic population in the BCL-X_L_-inhibitor sensitive CAKI-2 and A-498 cells, but not the relatively insensitive UMRC-6 and UMRC-2 cells (fig. S3). Together, these results demonstrated that pharmacological tools produced results that were largely consistent with our genetic studies and linked the dependency of ccRCCs on BCL-X_L_’s anti-apoptotic role.

We next attempted to validate these findings using an orthogonal assay. Anti-apoptotic BCL-2 family proteins exert their function by binding and sequestering pro-apoptotic “BH3-only” proteins (e.g. BIM, BID, and PUMA) or the pore-forming proteins (e.g. BAX and BAK). Synthetic, pro-apoptotic BH3 peptides can mimic or block these interactions and trigger cellular apoptotic programs. Probing for hallmarks of apoptosis, such as mitochondrial membrane depolarization that is indicative of cytochrome c release, in response to these peptide treatments can measure cellular dependencies on specific anti-apoptotic BCL-2-family proteins *ex vivo* (*29*). Using this “BH3 profiling” approach, which monitors mitochondrial potential using the JC1 dye, we compared the dependencies of ccRCCs on BCL-2-family anti-apoptotic proteins. In line with our pharmacological experiments, we observed that the BCL-X_L_ dependent CAKI-2 cells showed significant membrane depolarization with agents that blocked BCL-X_L_’s interaction with downstream effectors (e.g. the Hrk peptide and the Wehi539 pharmacological inhibitor), but not with MCL-1 blockers (MS1 peptide) (Fig. 2C). In contrast, the UMRC-2 cells, which were less responsive to acute BCL-X_L_ inhibition, were also largely non-responsive to agents that disrupt BCL-X_L_ function (Fig. 2D). These results demonstrated that BCL-X_L_ actively restrains pro-apoptotic signaling in ccRCCs, leading to a dependence on this protein for continued survival.

### BCL-X_L_ inhibition sensitizes kidney cells to chemotherapeutic agents

RCCs have historically been recognized for their resistance to traditional chemotherapeutics (*30*), most of which function by promoting apoptosis. We hypothesized that BCL-X_L_ function, which has been previously shown to be a drug resistance mechanism in many cancers, could be a physiological barrier to chemotherapeutic response in ccRCC. To address this hypothesis, we treated the A-1331852-insensitive UMRC-2 and OSRC2 cells with various dosage combinations of A-1331852 and chemotherapeutics such as 5-FluoroUracil (5-FU), Docetaxel, and Doxorubicin. Many statistical models exist to analyze drug-drug interaction matrices to score for potential synergy or antagonism (e.g. HSA, Loews, Bliss, and ZIP) (*31*–*34*). To overcome the caveats associated with these individual models, open source R-based packages have been generated, which simultaneously compare the dataset across multiple models. We used a recently developed we application that provides visualization of the SynergyFinder package (*35*). This analysis demonstrated synergy at many dose combinations between the BCL-X_L_ inhibitor and all three chemotherapy drugs using all the models, which we then aggregated to identify doses of potential synergy between A-1331852 and the corresponding chemotherapeutic agent (Fig. 3, fig. S4, and Table S2). Together, our drug-drug interaction studies showed that BCL-X_L_ inhibition chemosensitized ccRCCs to a wide range of chemotherapeutic agents.

**FIGURE 3.**
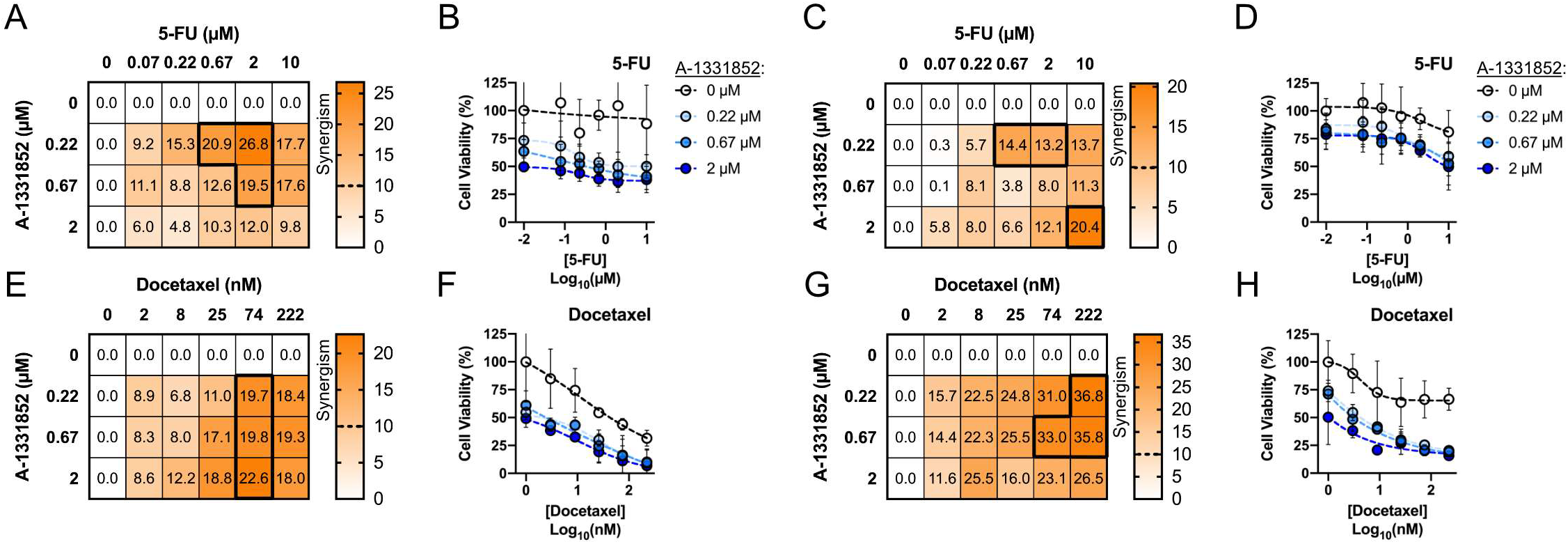
BCL-X_L_ Inhibition Chemosensitizes ccRCCs to Chemotherapeutics. (**A, C, E, and G**) Mean synergism score calculated across four independent models (ZIP, HSA, Loewe, Bliss) for 5-fluorouracil (5-FU) (**A** and **C**) and Docetaxel (**E** and **G**) at the indicated doses in combination with A-1331852. **(B, D, F, and H)** Percent cell viability and calculated dose-response curves in cells cultured in the presence of the indicated concentrations of the two molecules for 72 hours. Results for UMRC-2 (**A**, **B**, **E**, and **F**) and OSRC2 (**C**, **D**, **G**, and **H**) were calculated from ≥3 biological replicates. Error bars represent mean±SD.

### Epithelial-Mesenchymal Transition (EMT) is Associated with Bcl-xL Dependency

Our findings suggested that a subset of ~40% ccRCC lines were sensitive to acute BCL-X_L_ inhibition. Therefore, we interrogated the biomarkers that would allow differentiating ccRCCs based on BCL-X_L_ dependence. To this end, we first mined the Broad Achilles dataset and correlated mRNA levels of *BCL2L1* in different lineages to their respective BCL-X_L_ dependency using Pearson’s correlation coefficient. This analysis showed a weak correlation (r^2^ < 0.3) between *BCL2L1* mRNA expression levels and BCL-X_L_ dependency (fig. S5A). Next, we compared the protein expression levels of BCL-X_L_ and its closely related family member, BCL-2, in our panel of ccRCC lines. Consistent, with the mRNA correlation analysis, we did not find any significant difference in BCL-X_L_ protein expression levels among the A-1331852 sensitive versus insensitive ccRCC lines (fig. S5B). We then compared the relative abundance of BCL-2 and BCL-X_L_ and observed that BCL-X_L_:BCL-2 ratio was also not a predictor of BCL-X_L_ dependency in ccRCC (fig. S5, C to E).

To identify determinants of BCL-X_L_ dependency in an unbiased manner, we performed transcriptomics studies. We chose two representative sensitive lines (A-498 and CAKI-2) and two insensitive lines (OSRC-2 and UMRC-2). We transcriptionally profiled, using RNA-Seq, these ccRCC cells under (untreated) native growth conditions and also after acute treatment with A-1331852 (Fig. 4A). We reasoned that this approach would not only allow us to measure innate differences among cells that might underlie BCL-X_L_ dependence, but also potential differences in cellular response to acute treatment with BCL-X_L_ inhibitors. For these experiments, we empirically determined A-1331852 treatment time-points (typically 6-16 hour treatments) that preceded any overt cytotoxicity, which we feared could confound our gene expression studies.

**FIGURE 4.**
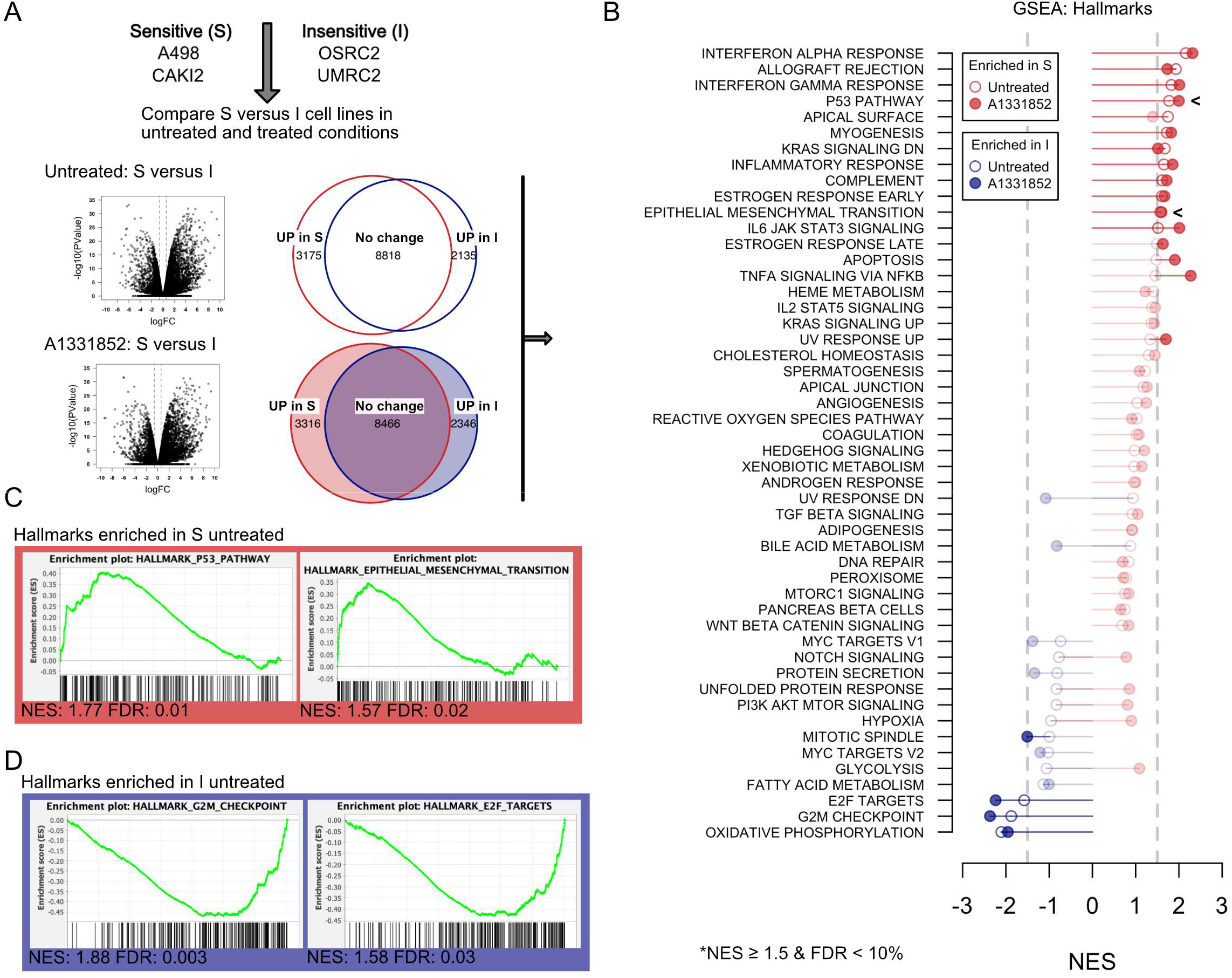
Transcriptomics Analysis Identifies Correlates Associated with BCL-X_L_ Dependency. (**A**) Schema (top) and number of genes that were identified as differentially expressed (bottom, ≥ 1.5-fold and FDR ≤ 10%, as determined by RNA-Seq) in the indicated BCL-X_L_ inhibitor ‘Sensitive’ (S) versus ‘Insensitive’ (I), under untreated basal conditions or upon acute treatment with A-1331852 (as described in the Methods). (**B** and **C**) Lollipop plot (**B**) depicting statistically significant results (NES ≥ 1.5 and FDR ≤ 10%) of Gene Set Enrichment Analysis comparing RNA-Seq data from S vs I cells that were either untreated (open circles) or A-1331852 treated (filled circles) and representative gene sets that were enriched in the Sensitive cells (top, red) (**C**) or the Insensitive cells (bottom, blue) (**D**).

We analyzed the transcriptional signatures using the EdgeR module in R and identified genes that were differentially expressed among the sensitive (S) versus insensitive (I) lines in both the untreated and the A-1331852-treated condition (Fig. 4A and Tables, S3 and S4). Among these differentially expressed genes, we compared enrichment or depletion of annotated gene sets described in mSigDB (*36*), using Gene Set Enrichment Analysis (*37*). These studies found that, compared to the insensitive cells, prominent differences existed innately in untreated sensitive cells (Fig. 4B and Table. S5); however, the extent of many these differences were further amplified upon BCL-X_L_ inhibition (Fig. 4B and Table. S6). These gene sets represented a number of biological pathways, including p53 response, cytokine response (e.g. interferon response), and cell state pathways [e.g. epithelial to mesenchymal transition (EMT)] (Fig. 4, B to D, and Tables, S5 and S6).

When activated by cellular damage or stress, p53 transcribes multiple pro-apoptotic proteins to trigger cell death in cells that are primed for apoptosis (*38*, *39*), and favors cell death induction by inhibitors of BCL-X_L_ (*40*). We therefore reasoned that intact p53 signaling may be associated with BCL-X_L_ dependence and potentially act as a predictor of A-1331852 response. We thus tested whether the segregation of ccRCC lines either into BCL-X_L_-dependent or independent subsets was p53-dependent. First, we mined the Broad Institute’s Cancer Cell Line Encyclopedia (CCLE) to evaluate p53 mutation status in our panel of ccRCC lines. This analysis showed that 3 of the 4 sensitive cell lines had wild-type p53; however, the p53-mutant SLR23 cells were also highly sensitive to BCL-X_L_ loss (fig. S6A). In contrast, cell lines that were insensitive to BCL-X_L_ inhibition included OSRC2 and UOK101 cells, both of which have wild-type p53 (fig. S6A). Therefore, p53 status alone appeared to be insufficient to predict BCL-X_L_ dependency.

To probe this further, we treated ccRCC lines with doxorubicin and measured the accumulation of total p53, phosphorylated p53, and the p53-target gene *CDKN1A*, which encodes the cyclin-dependent kinase inhibitor p21. We found that the basal levels of p21, and its induction in response to doxorubicin, was lower in many of the intermediate and insensitive ccRCC lines, compared with the sensitive cells (fig. S6B). UOK101 cells were, once again, a notable exception with significant p21 expression but little BCL-X_L_ dependence, perhaps due to other mutations that block apoptosis in this cell line. Altogether, we concluded that p53 response to DNA damaging agents could partly predict BCL-X_L_ dependency in ccRCC lines.

Finally, we focused our attention on the cell state pathways because of the relative novelty of epithelial-mesenchymal transition (EMT) as a determinant of BCL-X_L_ dependency. To validate our RNA-Seq data, we used flow cytometry to compare the expression levels of CD44, a well characterized marker of mesenchymal state, in ccRCC cells that were either sensitive or insensitive to acute BCL-X_L_ inhibition. Consistent with our transcriptional studies, we observed that CD44 expression levels were significantly elevated in BCL-X_L_ dependent cells (fig. S7A). These studies suggested that mesenchymal tumor cell state was indeed a promising biomarker of BCL-X_L_ dependence.

We then addressed the functional causality of this pathway in driving apoptosis upon A-1331852 treatment. We reasoned that a causal role for cell state differences in conferring BCL-X_L_ dependence could be established if downregulating mesenchymal features in sensitive cells promoted insensitivity to A-1331852; or conversely, if inducing mesenchymal features in insensitive lines was sufficient to sensitize them to A-1331852 treatment. To this end, we first treated the sensitive lines, A-498 and CAKI-2, with all-trans-Retinoic Acid (ATRA) for three days with the hope that the differentiation promoting effects of ATRA would diminish mesenchymal features in these cells (Fig. 5A). Indeed, we noted a small, but statistically significant, reduction in CD44 expression in both cell lines (Fig. 5B). Interestingly, this minimal change in mesenchymal was still sufficient to reduce the sensitivity of both A-498 and CAKI-2 cells to A-1331852 treatment (Fig. 5C).

**FIGURE 5.**
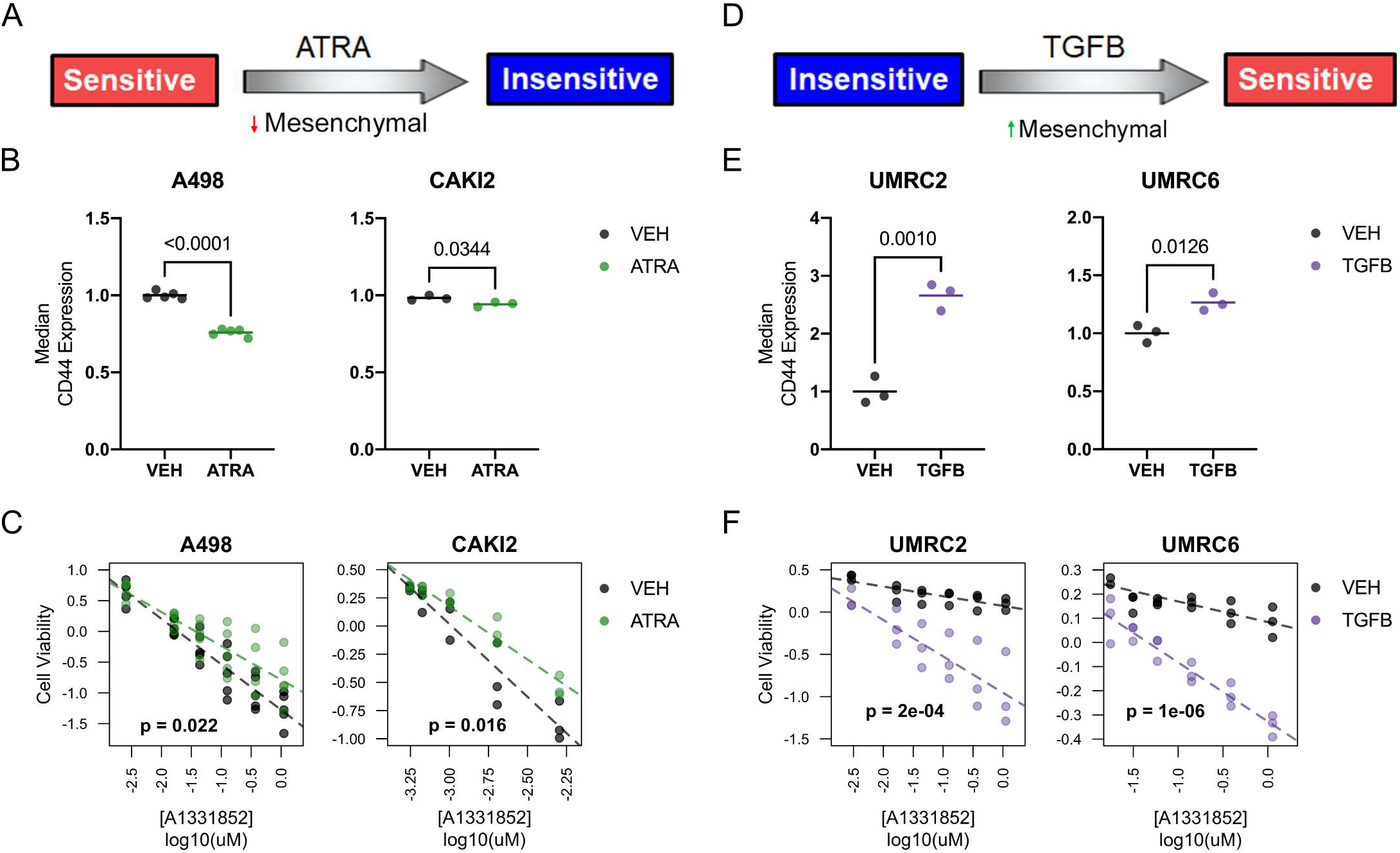
A Mesenchymal Cell State Promotes BCL-X_L_ Dependency. (**A**) Schema showing experimental design to address necessity of mesenchymal state in promoting BCL-X_L_ dependency. (**B**) CD44 levels, as determined by flow cytometry, in the indicated ccRCC cells that were treated with 1 μM ATRA for 3 days. (**C**) Cell viability, relative to untreated DMSO controls, in the indicated cells treated with A-1331852 for 7 days. (**D**) Schema showing experimental design to address sufficiency of mesenchymal state in promoting Bcl-xL dependency. (**E**) CD44 levels, as determined by flow cytometry, in the indicated ccRCC cells that were treated with 10 ng/ml TGFβ for 3 days. (**F**) Cell viability, relative to untreated DMSO controls, in the indicated cells treated with A-1331852 for 7 days. Data in (**B**) and (**E**) was normalized to CD44 levels in the untreated (Vehicle) control and was compared using the Student’s t-test (n ≥ 3, bar represents Mean, p-values are indicated). (**C**) and (**F**) were normalized for batch effects and compared using linear regression (n ≥ 3, dotted line represents best fit, p-values indicated are for difference in slopes i.e., interaction between A-1331852 and ATRA or TGFβ)

Next, we designed an experiment to address mesenchymal sufficiency. We began by treating the A-1331852-insensitive, UMRC-2 and OSRC-2, cells for 3 days with TGFβ, a potent inducer of mesenchymal transition (Fig. 5D). Unfortunately, we did not find any changes in CD44 expression in OSRC2 cells and thus substituted these cells with the intermediate-sensitive UMRC-6 cells. In both UMRC-2 and UMRC-6 cells, TGFβ treatment led to robust induction of CD44 (Fig. 5E). Strikingly, promoting mesenchymal features also increased A-1331852 sensitivity in both these lines (Fig. 5F). Importantly, these results were specific to BCL-X_L_ dependence because perturbation of cell state failed to confer sensitivity to pharmacological BCL-2 inhibition (fig. S7, B and C). Altogether, these studies demonstrated that a mesenchymal signature did not merely correlate with increased BCL-X_L_ dependency but was both necessary and sufficient to modulate the response of ccRCC cells to BCL-X_L_ blockers.

Previous literature suggested that chronic HIF activation in pVHL-deficient ccRCCs status could impact apoptotic response. Many transcriptional targets of HIF are known regulators of cell death pathways, including *BCL2L1* in certain cellular contexts (*41*, *42*). Moreover, in certain cell lines, pVHL expression has been shown to impact EMT (*43*, *44*). Therefore, we addressed if pVHL status could influence BCL-X_L_ dependency. We lentivirally transduced pVHL-deficient ccRCC cells to restore expression of wild-type pVHL (VHL) or an empty vector control (VEC), and thus generated isogenic CAKI-2 and A-498 cells that either did or did not express pVHL (Fig. S8A). Reintroduction of pVHL does not lead to fitness defects *in vitro* under standard cell culture conditions, but impedes ccRCC tumor growth *in vivo*, enabling *in vitro* studies that address the influence of pVHL status. Surprisingly, reintroduction of pVHL in CAKI2 and A498 cells did not alter their mesenchymal cell state (fig. S8B), and consistent with these observations failed to impact the sensitivity of these cells to pharmacological BCL-X_L_ inhibition (fig. S8, C and D). These results indicate that the impact of pVHL on EMT is likely dependent on biological context, and pVHL status is not sufficient to predict BCL-X_L_ dependence in kidney cancer cells.

### The EMT Signature Predicting BCL-X_L_ Dependence is a Clinically-exploitable Feature

Based on the importance of the cell state pathways as determinants of BCL-X_L_ dependency, we next probed for the prevalence of this signature in clinical samples. We generated a list of differentially-expressed genes (DEG) by comparing the relative expression patterns between the A-1331852-sensitive and insensitive ccRCC cell lines (Figs. 4A and 6A). We then analyzed kidney cancer gene expression data mined from The Cancer Genome Atlas (TCGA) and noted, as expected, that principal component analysis led to segregation of the RCC tumors into three distinct clusters, each representing one major RCC disease subtype (Fig. 6B and fig. S9). We then integrated the DEG list predicting BCL-X_L_ dependence with the ccRCC tumor cluster. This analysis allowed us to establish (a) the prevalence of the BCL-X_L_ dependency signature (Fig. 6C), and (b) the differences in amplitude of the individual gene sets that together comprised the BCL-X_L_ dependency signature, specifically in human clear cell renal tumors. We observed that the BCL-X_L_ dependency signature (red) was observed in ~30% of human ccRCCs. The EMT signature was among the most differentially expressed gene signatures among these cohorts; whereas, differences in the p53 pathway, albeit statistically significant, showed more subtle differences (Fig. 6D). Finally, as expected from the typically aggressive nature of mesenchymal tumors, the BCL-X_L_ dependency signature was associated with worse clinical outcomes (Fig. 6E). These studies suggested that mRNA gene expression could be exploited as a biomarker to identify patient tumors exhibiting features of BCL-X_L_ dependence.

**FIGURE 6.**
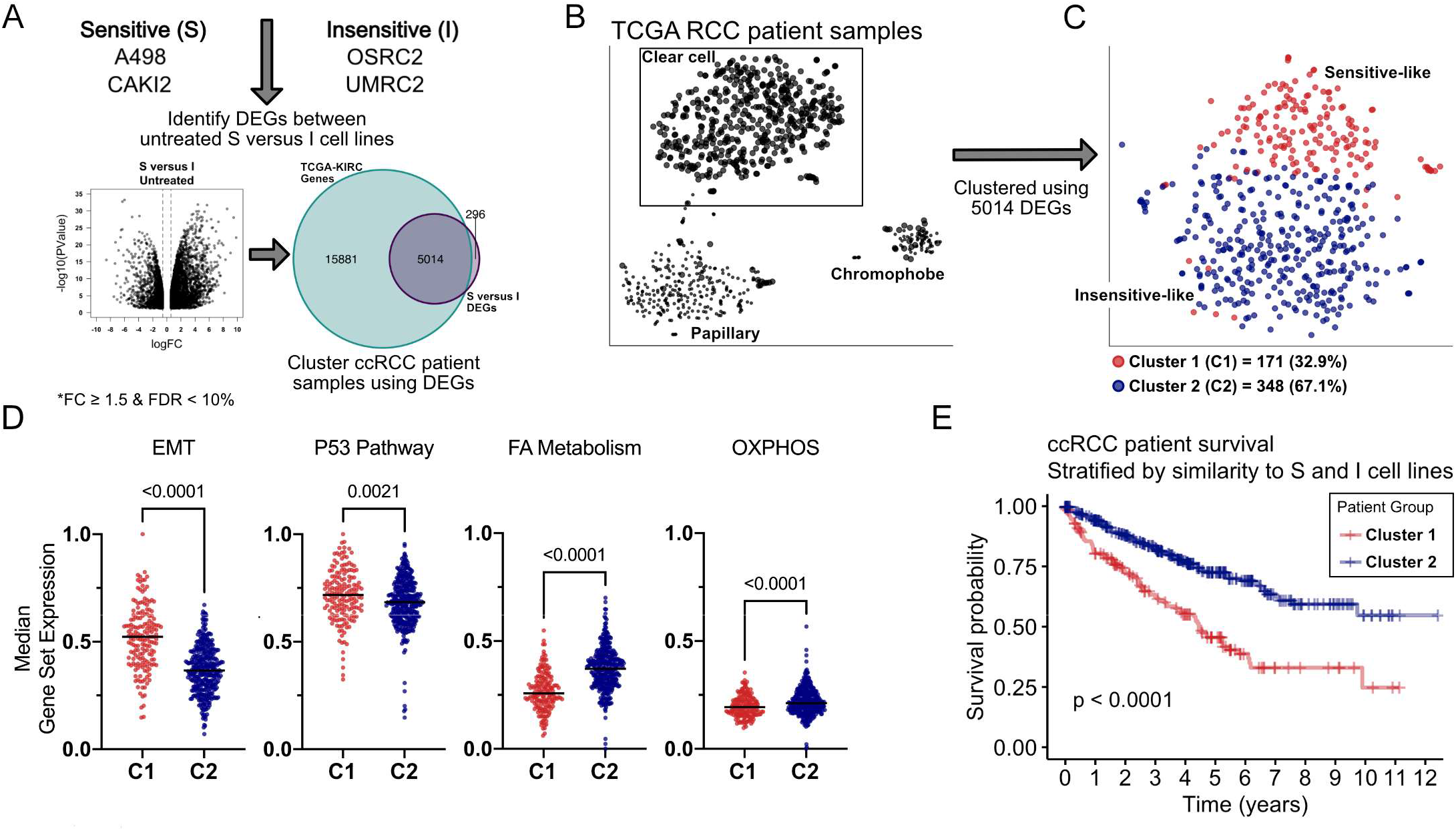
BCL-X_L_ Dependency Signatures are Evident in ccRCC Clinical Specimens. (**A**) Schema indicating the computational overlay of differentially-expressed genes, as determined in 4A, onto renal cancer gene expression data mined from TCGA. (**B**) Principle-component analysis showing segregation of tumors based on their subtype (left), and then within ccRCC (**C**) by similarity in gene signature to Sensitive (red) versus insensitive (blue) lines (right). (**D** and **E**) Median gene expression levels of the indicated gene sets (**D**) and Kaplan-Meier curves depicting clinical outcomes (**E**) in patients with tumors resembling the BCL-X_L_ inhibitor Sensitive versus Insensitive cell lines. (**A**) and (**D**) were compared using the Student’s *t-test*. In (**E**) Sensitive (n = 171) and Insensitive (n = 348) patients were compared by the log-rank test.

### BCL-X_L_ Inhibition Impedes In Vivo Tumor Growth

The BCL-XL-selective inhibitor, A-1331852, has been previously shown to be orally bioavailable in subcutaneous tumor xenografts (*45*). Therefore, A-1331852 treatment could be used to interrogate the therapeutic feasibility of BCL-X_L_ inhibition in ccRCC. Unfortunately, several ccRCC lines, including the highly BCL-X_L_-dependent CAKI-2 and A-498 cells, have poor engraftment rates as subcutaneous tumors in NCR^nu/nu^ mice. The UMRC-2 cells, however, engraft tumors readily. Interestingly, we found that, as seen with our genetic studies, sustained pharmacological BCL-X_L_ inhibition (>2 weeks) led to nM IC_50_ in UMRC-2 cells (fig. S10). These results justified the use of UMRC-2 cells for *in vivo* studies, which typically rely on dosing regimens extending to 3-4 weeks.

We inoculated UMRC-2 cells subcutaneously into 7 week old NCR^nu/nu^ mice (equal number of males and females) and allowed tumor engraftment and growth for ~6-8 weeks. Tumor-bearing animals (>100 mm^3^) were randomized to receive either 25 mg/kg A-1331852 or sham-vehicle control, twice a day, orally, for up to 4 weeks. Tumor volumes were measured at the point of enrollment (Fig. 7A) and then weekly after dosing was initiated (Fig. 7B). We noted a significant decrease in tumor growth in A-1331852 treated mice, as noted in smaller tumor volumes (Fig. 7, B to D). Histological analysis of the harvested tumors showed zones of necrotic cells in A-1331852 treated tumors. These necrotic regions were marked by elevated Cleaved Caspase 3 staining, consistent with apoptotic cell death triggered by A-1331852 treatment (Fig. 7, E and F).

**Figure 7.**
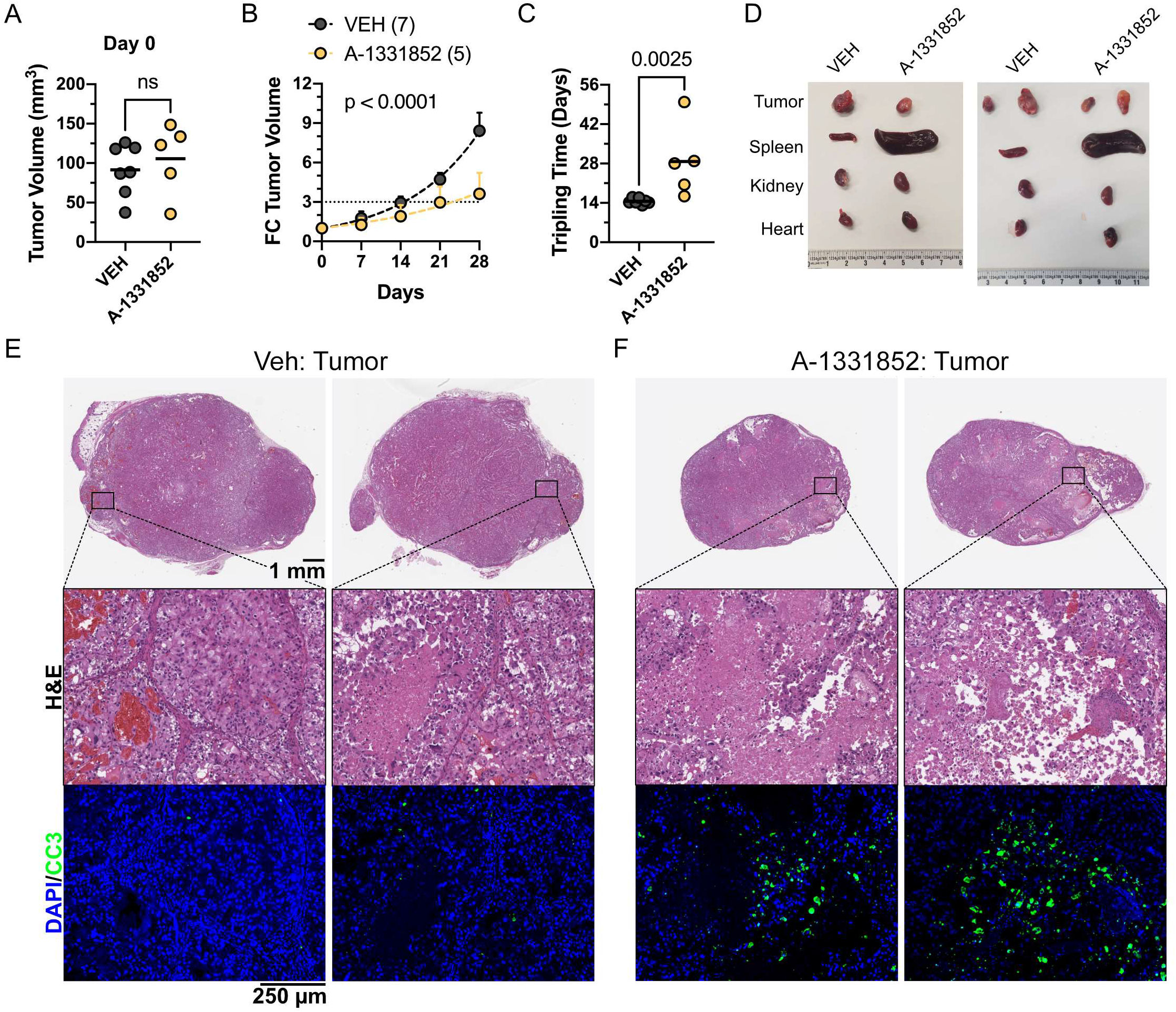
Pharmacological BCL-X_L_ Inhibition Impedes Tumor Growth. Tumor volumes at Day 0, start of dosing (**A**), change in tumor volume at the indicated time points after A-1331852 dosing at 25 mg/kg, twice a day, by oral gavage (**B**), time for tumor volume to triple on the indicated dosing regimens (**C**), and photomicrographs of harvested tumors and the indicated animal organs (**D**), in NOD/SCID mice that were inoculated subcutaneously with pVHL-deficient UMRC-2 cells. (E and F) Histological analysis by H&E staining and immunohistochemistry, showing DAPI (blue) and Cleaved Caspase 3 (green) staining (bottom panel), of tumors harvested from animals dosed with Vehicle control (**E**) or A-1331852 (**F**), as indicated.

The animals handled the dosing without any notable behavioral or body weight changes (fig. S11A). However, we noted profound splenomegaly in A-1331852 treated mice (Fig. 7D and fig. S11B). Histological analysis confirmed a notable disruption of splenic tissue architecture in A-1331852-treated animals, including a virtual absence of germinal centers (fig. S11, C and D). In contrast to the spleen, however, the histology and macroscopic appearance of other organs (e.g. kidney, liver, lung, and heart) were virtually indistinguishable between the vehicle and A-1331852 treated cohorts (fig. S12). Altogether, these findings were promising despite the technical necessity to employ the relatively insensitive UMRC-2 cells, and demonstrated the overall feasibility of BCL-X_L_ inhibition as a strategy to block ccRCC tumor growth.

## DISCUSSION

Despite recent advances in therapy, metastatic kidney cancer is ultimately incurable, and there is a recognized need for new therapeutic approaches to complement existing treatments. To address this need, in this study we interrogated lineage-specific dependencies. We made two modifications to a standard analytical pipeline. First, rather than probing for ‘synthetic lethality’ dependencies that are associated with a specific oncogenic alteration (e.g. truncal genetic lesion), we mined for candidate dependencies that are preferentially enriched in a given lineage, here in kidney cancer. Second, within the lineage, we sorted for – and retained – strongly selective dependencies even in a subset of cells. This strategy enabled the identification of candidate “hits” that were strong subset-specific dependencies in kidney cancer cells, including *BCL2L1*, which encodes the anti-apoptotic BCL-X_L_ protein.

We focused on BCL-X_L_ because it was both druggable and already had many well characterized pharmacological inhibitors available, which could facilitate faster adaptation into the clinic. However, other genes in our list (e.g. *ITGAV*) also encode druggable proteins and warrant further study. Moreover, some otherwise “undruggable” proteins (e.g. HNF1B and PAX8) could be targeted with drugs that promote their proteolysis, such as deubiquitinase inhibitors and/or proteolysis-targeting chimeras (PROTACs). Notably, PROTACs have also been developed for BCL-X_L_, in an attempt to reduce drug resistance and to reprogram BCL-X_L_ inhibition away from platelets, where it leads to undesirable cytotoxic side effects.

We validated BCL-X_L_ as a dependency in ccRCC multiple ways. First, we confirmed that ccRCC lines, such as CAKI-2 and TUHR4TKB, which were exceptionally sensitive to *BCL2L1* in the Achilles dataset, were also independently identified as BCL-X_L_ dependent in the Novartis DRIVE genome-wide shRNA screen and the Broad Institute’s CRISPR/Cas9 dependency map. Next, using three different shRNAs, we confirmed fitness defects in multiple ccRCC cell lines upon loss of BCL-X_L_, but not its closely related sibling BCL-2. Finally, we demonstrated that these genetic effects were “on-target” by rescuing the cytotoxicity associated with two of these shRNAs.

To develop upon our genetic studies, we employed pharmacological agents that target BCL-X_L_. We found remarkable overlap in the sensitivity of cell lines using either one of these approaches. We also noted that some cell lines (e.g. UMRC-2) showed cytotoxicity only upon chronic BCL-X_L_ blockade. Importantly, we did not observe any measurable response to the BCL-2 inhibitors, demonstrating the specificity of BCL-X_L_ dependence in ccRCC.

Kidney cancer, until recently, was curable only by surgical resection. This was largely because standard chemotherapeutic agents, including topoisomerase inhibitors, taxanes, and nucleoside analogues failed to show any significant clinical efficacy in many independent trials (*30*). The ability to promote apoptosis is central to the efficacy of many traditional chemotherapeutic agents. We therefore addressed if the anti-apoptotic function of BCL-X_L_ could be a physiological barrier to chemotherapy response, and thus underlie the observed insensitivity of ccRCCs to these agents. Consistent with this idea, our drug-drug interaction studies showed synergistic interactions between the BCL-X_L_ inhibitor A-1331852 and a number of traditional chemotherapeutics. Therefore, BCL-X_L_ inhibition sensitizes RCCs to chemotherapy, and offers us clinically usable opportunities to target RCCs that are non-responsive to other therapeutic strategies.

The importance of anti-apoptotic proteins has been previously interrogated in renal cancer; however, these studies focused primarily on the role of BCL-2 because BCL-2 inhibition had been a successful strategy to target other cancers (*46–48*). Our findings, relying on unbiased screens and subsequent validations, demonstrated that kidney cancer cells exhibit much greater dependence on BCL-X_L_ than BCL-2. These findings refocus attention on BCL-X_L_ blockers as potential therapeutic agents in kidney cancer.

Our findings indicate that ~30% of human renal tumors, especially those that represent the more aggressive mesenchymal signatures, are likely to be responsive to Bcl-xL inhibition. To further this idea, we report a number of biomarkers that can be measured in tumor biopsies, including p21 expression (as a marker for p53 activity) and mesenchymal mRNA gene expression signatures. We also demonstrate the utility of the *ex vivo* BH3-profiling assay as a faithful predictor of BCL-X_L_ dependency in renal cells. Importantly, this assay has been optimized for interrogation of apoptotic dependencies in human cancer samples (*29*, *49*), and can be easily adapted for human renal tumors. Together, these assays provide us numerous future opportunities, which were beyond the scope of this initial study, to develop clinically usable biomarkers to identify cohorts of BCL-X_L_ dependent renal tumors.

The clinical use of BCL-X_L_ blockers has been limited due to adverse events, raising concerns about the potential clinical utility of this approach. However, most of these studies were performed using earlier versions of BCL-X_L_ blockers, which also had effects on BCL-2 and other BCL-2-family proteins including BCL-w. More specific BCL-X_L_ inhibitors (such as the highly potent agent A-1331852) may have reduced toxicities when they are evaluated *in vivo*. Thrombocytopenia will continue to be a concern given that platelets are highly dependent on BCL-X_L_ for survival and quickly undergo apoptosis when this protein is inhibited (*50*). However, novel approaches that degrade BCL-X_L_ in nucleated cells but not platelets reduce thrombocytopenia in animal models and are now entering clinical trials (*51*). These pharmacological advances, combined with assays that exploit our biological predictors of BCL-X_L_ dependence and assign therapy more rationally in patients most likely to respond, could enable efficacious clinical use of BCL-X_L_ inhibitors in kidney cancer.

Why might mesenchymal cancer cells be dependent on BCL-X_L_? In normal tissues, cellular shedding in response to disruption of epithelial organization triggers cell death pathways in dislodged cells. This process is called Anoikis (greek for “homelessness”) [fig. S13 (left)]. Cancer cells undergoing epithelial-mesenchymal transition, should likewise be primed for cell death. We speculate that mesenchymal kidney cancer cells possibly begin to resemble an anoikis-like state and select for BCL-X_L_’s anti-apoptotic activity to ensure their survival [fig. S13 (right)]. Altogether, our observations justify testing the use of BCL-X_L_ blockers in the management of the mesenchymal – more aggressive – kidney tumors.

## Acknowledgments

We thank Dr. Ruth Keri (Cleveland Clinic) for the BCL-XL expression construct and Dr. Alexandru Almasan (Cleveland Clinic) for the SU-DHL-6 cell line. We thank Dr. Christopher Weight (Cleveland Clinic) for critiques on the clinical relevance of these findings.

## Author contributions

SM and LS designed and performed cell-based experiments. TR modeled the drug-drug interactions. CF and KAS performed BH3 profiling studies. JMKB and FV analyzed the Achilles data to generate the candidate gene list. TG, AAC, and WGK designed experiments, analyzed results, and wrote the manuscript.

## Data and materials availability

All gene expression data are deposited into the Gene Expression Omnibus (GSE173618).

