## Supplemental Figures for "A Mesenchymal Tumor Cell State Confers Increased Dependency on the BCL-X_L_ Anti-apoptotic Protein in Kidney Cancer"

FIGURE S1

A

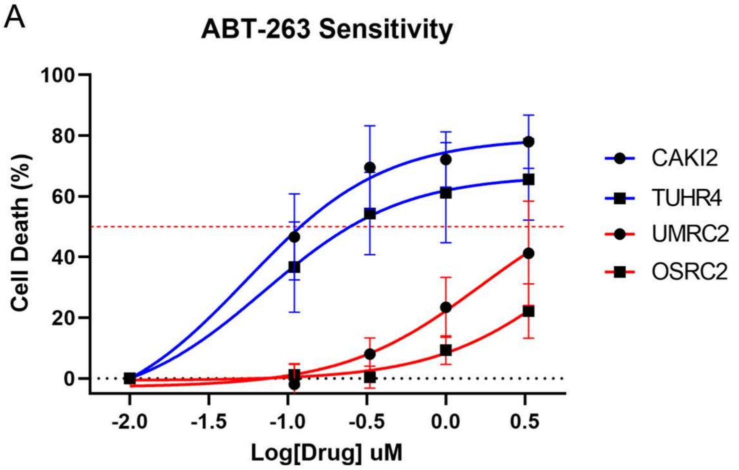


B

| Cell Line | $IC_{50}$ (μM) |
|---|---|
| CAKI2 | 0.05450 |
| TUHR4 | 0.07065 |
| UMRC2 | 1.629 * |
| OSRC2 | 6.457 * |

Figure S2

A

| Cell Line | A-1331852 $IC_{50}$ (µM) |
|---|---|
| CAKI2 | 2.17E-06 |
| A498 | 4.44E-05 |
| TUHR4 | 3.61E-06 |
| SLR23 | 0.01187 |
| 786O | ND |
| UMRC6 | ND |
| UMRC2 | ND |
| UOK101 | ND |
| OSRC2 | ND |
| RCC4 | ND |

B

| Cell Line | ABT-199 $IC_{50}$ (µM) |
|---|---|
| SUDHL-6 | 2.24E-06 |
| CAKI2 | ND |
| TUHR4 | ND |
| UMRC2 | ND |
| OSRC2 | ND |
| A498 | ND |
| 786O | ND |
| UOK101 | ND |
| RCC4 | ND |
| SLR23 | ND |
| UMRC6 | ND |

FIGURE S3

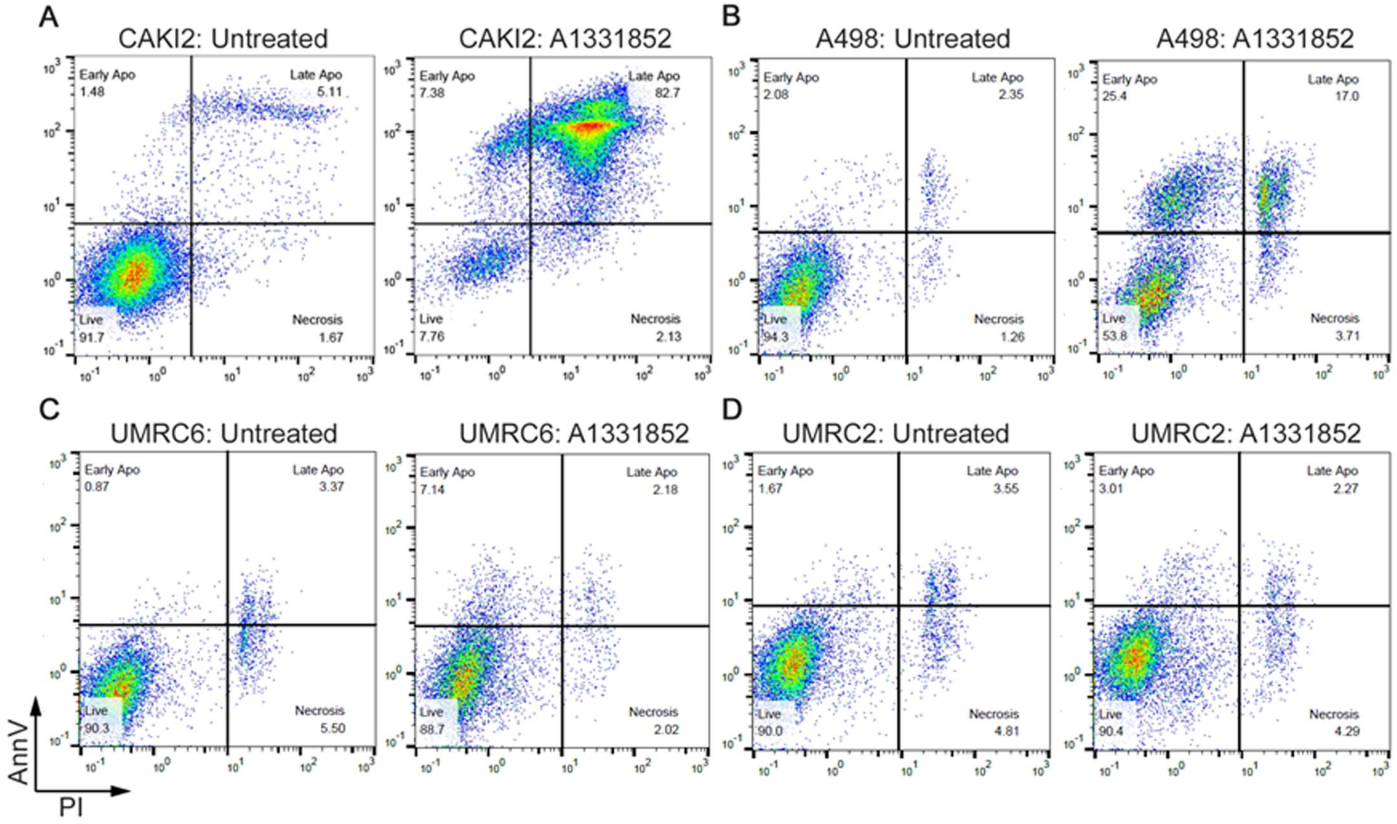
A
CAKI2: Untreated
Early Apo 1.48
Late Apo 5.11
Live 91.7
Necrosis 1.67
CAKI2: A1331852
Early Apo 7.38
Late Apo 82.7
Live 7.76
Necrosis 2.13
B
A498: Untreated
Early Apo 2.08
Late Apo 2.35
Live 94.3
Necrosis 1.26
A498: A1331852
Early Apo 25.4
Late Apo 17.0
Live 53.8
Necrosis 3.71
C
UMRC6: Untreated
Early Apo 0.87
Late Apo 3.37
Live 90.3
Necrosis 5.50
UMRC6: A1331852
Early Apo 7.14
Late Apo 2.18
Live 88.7
Necrosis 2.02
D
UMRC2: Untreated
Early Apo 1.67
Late Apo 3.55
Live 90.0
Necrosis 4.81
UMRC2: A1331852
Early Apo 3.01
Late Apo 2.27
Live 90.4
Necrosis 4.29
AnnV
PI

### FIGURE S4

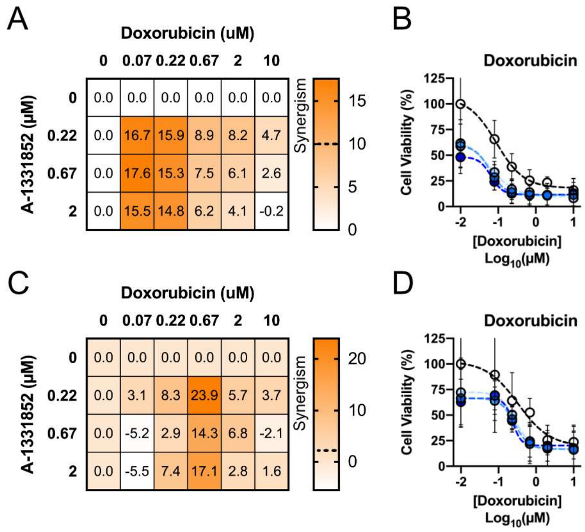
A
Doxorubicin (uM)
0 0.07 0.22 0.67 2 10
A-1331852 (µM)
0 0.0 0.0 0.0 0.0 0.0 0.0
0.22 0.0 16.7 15.9 8.9 8.2 4.7
0.67 0.0 17.6 15.3 7.5 6.1 2.6
2 0.0 15.5 14.8 6.2 4.1 -0.2
Synergism
15
10
5
0
B
Doxorubicin
Cell Viability (%)
125
100
75
50
25
0
-2 -1 0 1
[Doxorubicin]
Log10(µM)
C
Doxorubicin (uM)
0 0.07 0.22 0.67 2 10
A-1331852 (µM)
0 0.0 0.0 0.0 0.0 0.0 0.0
0.22 0.0 3.1 8.3 23.9 5.7 3.7
0.67 0.0 -5.2 2.9 14.3 6.8 -2.1
2 0.0 -5.5 7.4 17.1 2.8 1.6
Synergism
20
10
0
D
Doxorubicin
Cell Viability (%)
125
100
75
50
25
0
-2 -1 0 1
[Doxorubicin]
Log10(µM)

FIGURE S5

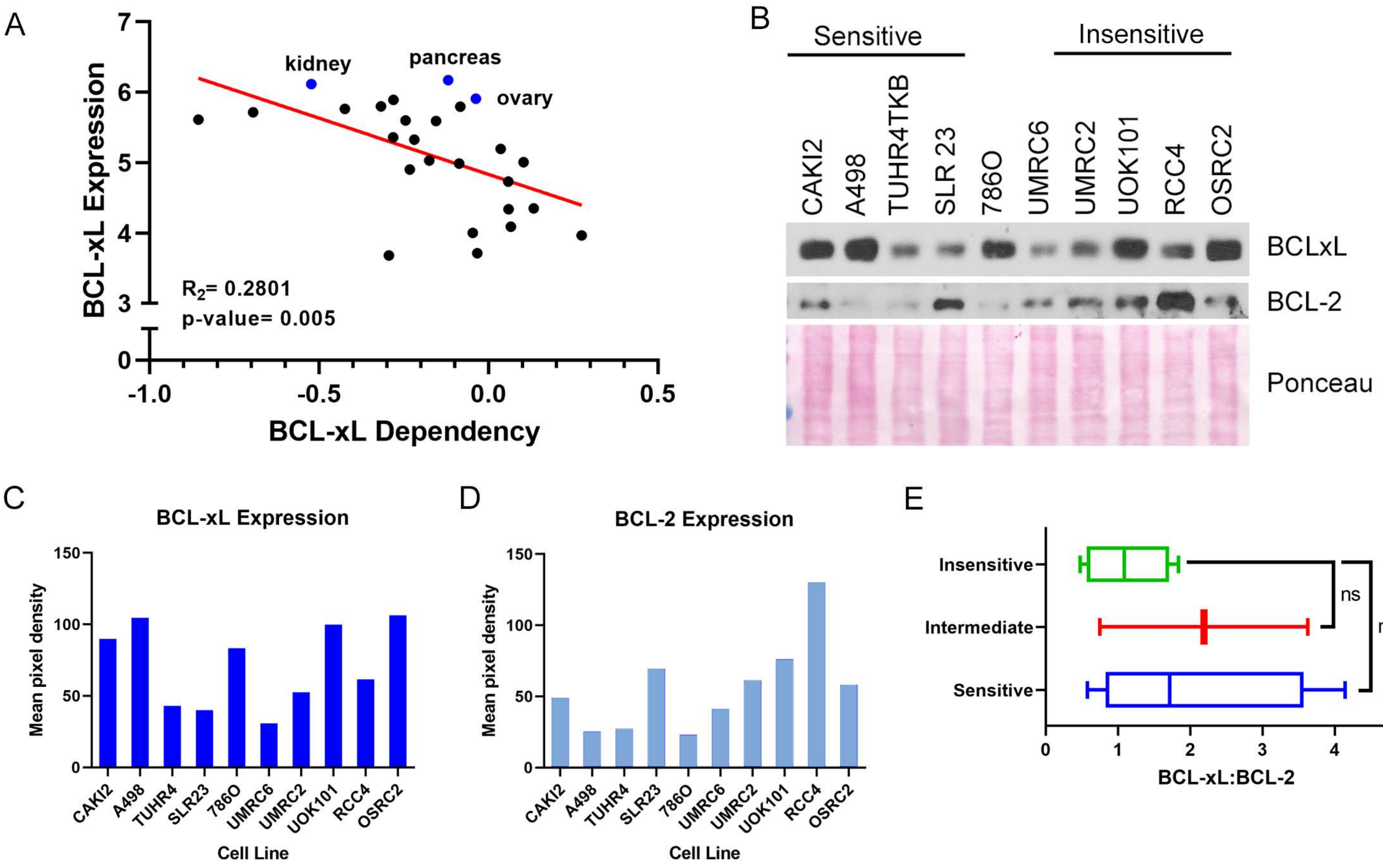
A
BCL-xL Expression
BCL-xL Dependency
kidney
pancreas
ovary
R2= 0.2801
p-value= 0.005
B
Sensitive
Insensitive
CAKI2
A498
TUHR4TKB
SLR 23
786O
UMRC6
UMRC2
UOK101
RCC4
OSRC2
BCLxL
BCL-2
Ponceau
C
BCL-xL Expression
Mean pixel density
Cell Line
D
BCL-2 Expression
Mean pixel density
Cell Line
E
Insensitive
Intermediate
Sensitive
ns
ns
BCL-xL:BCL-2

FIGURE S6

A

| | BCL-xL dep. | TP53 | VHL |
|---|---|---|---|
| CAKI2 | HIGH | Wild-type | R177* |
| A498 | HIGH | Wild-type | VD142fs |
| TUHR4TKB | HIGH | Wild-type | Y98fs |
| SLR23 | HIGH | R158H | W88* |
| 786O | INTERMED. | R248W | G104fs |
| | | R278A | |
| | | Splice Site | |
| UMRC6 | INTERMED. | | |
| UOK101 | LOW | Wild Type | T157fs |
| RCC4RGB | LOW | | |
| UMRC2 | LOW | | |
| OSRC2 | LOW | Wild Type | R58fs |

B

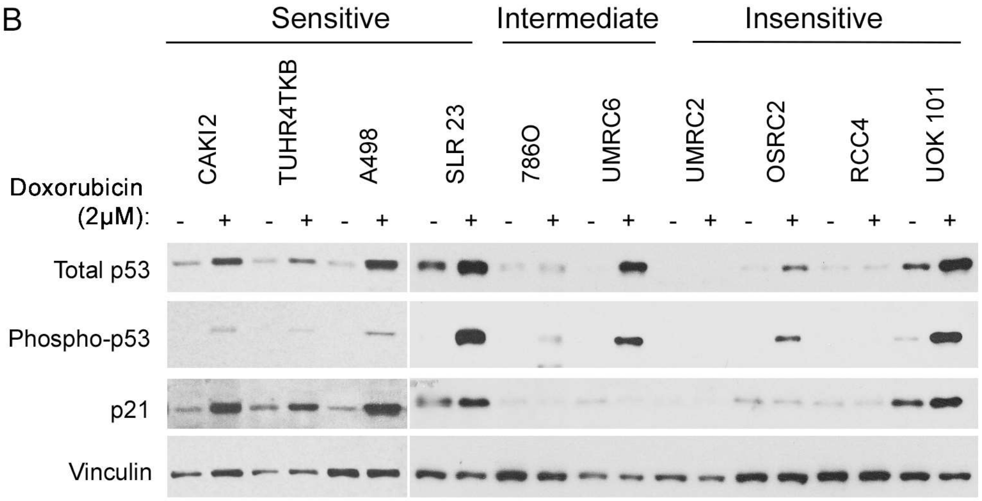

FIGURE S7

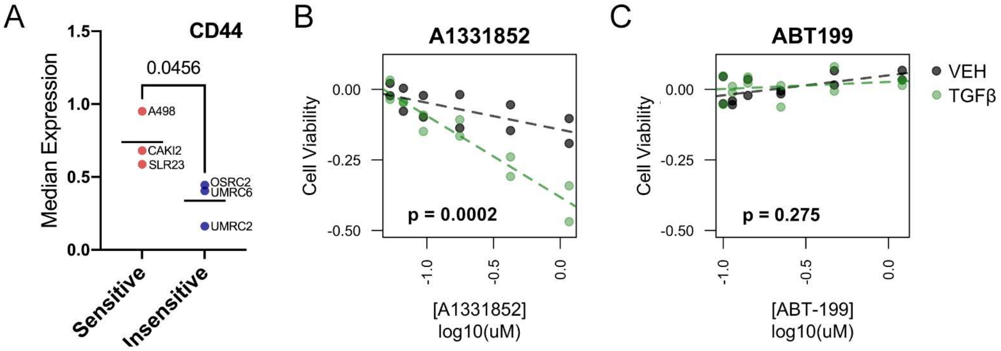
A
CD44
Median Expression
0.0456
A498
CAKI2
SLR23
OSRC2
UMRC6
UMRC2
Sensitive
Insensitive
B
A1331852
Cell Viability
p = 0.0002
[A1331852]
log10(uM)
C
ABT199
Cell Viability
p = 0.275
[ABT-199]
log10(uM)
VEH
TGFβ

FIGURE S8

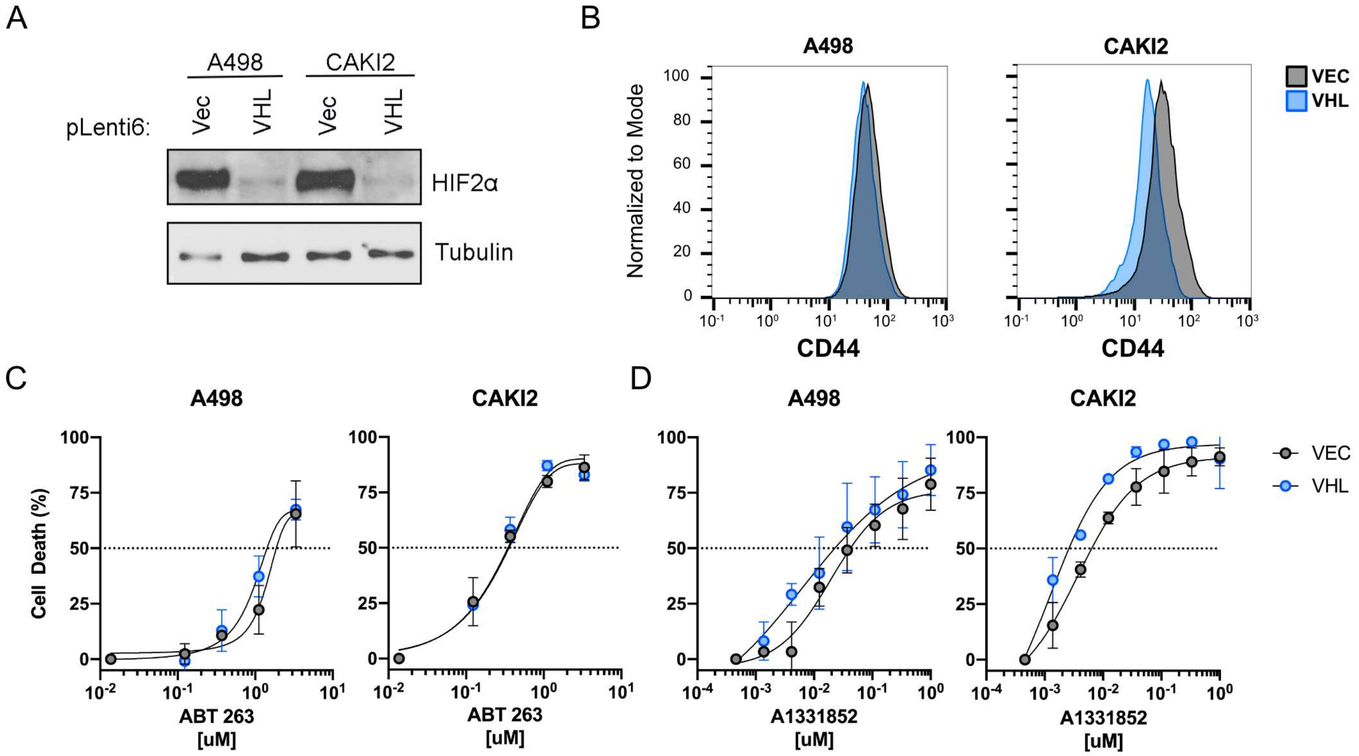
A
A498
CAKI2
pLenti6:
Vec
VHL
Vec
VHL
HIF2α
Tubulin
B
A498
CAKI2
VEC
VHL
Normalized to Mode
CD44
CD44
C
A498
CAKI2
Cell Death (%)
ABT 263
[uM]
ABT 263
[uM]
D
A498
CAKI2
VEC
VHL
A1331852
[uM]
A1331852
[uM]

FIGURE S9

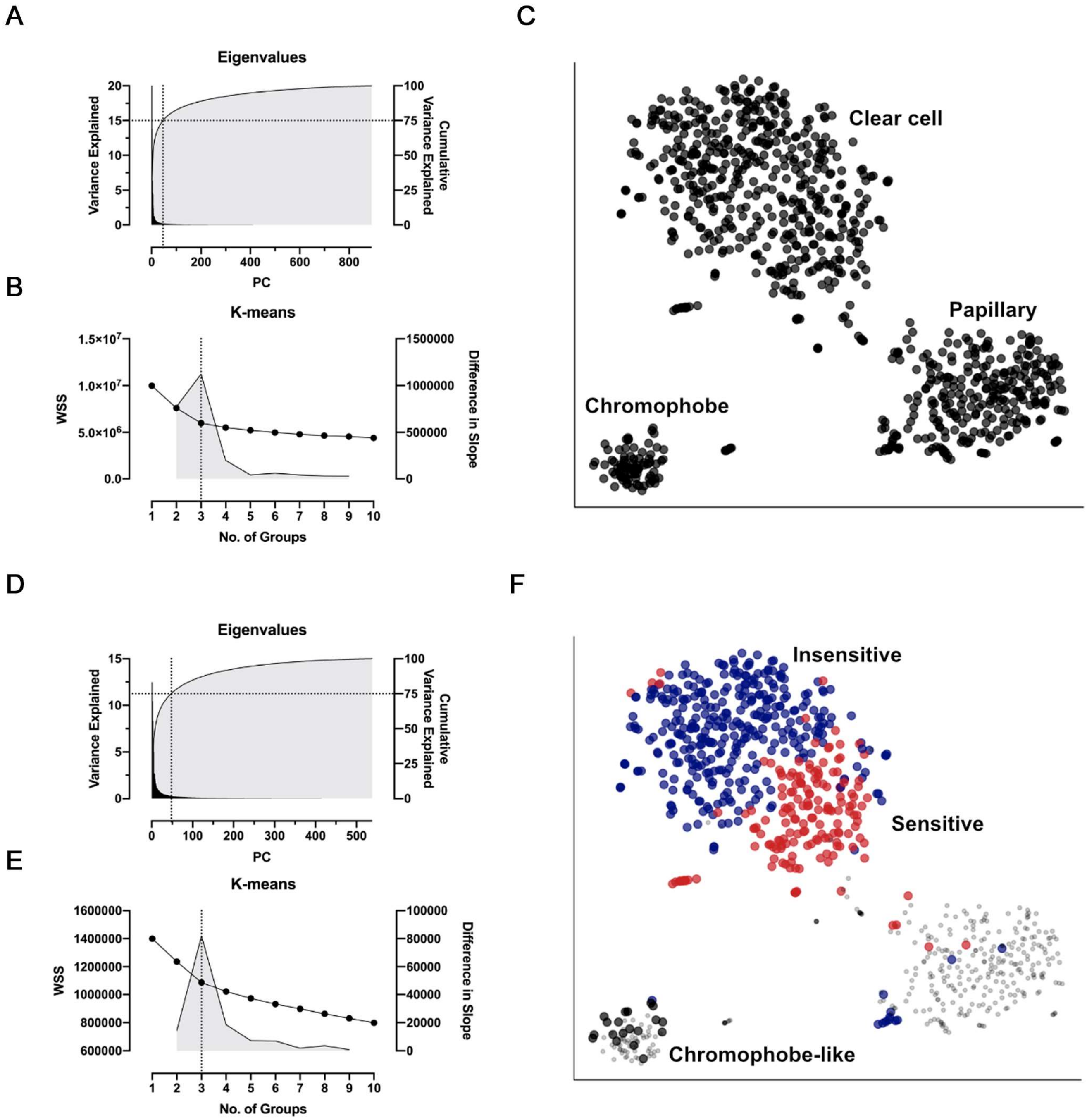
A
Eigenvalues
Variance Explained
Cumulative Variance Explained
PC
B
K-means
WSS
Difference in Slope
No. of Groups
C
Clear cell
Papillary
Chromophobe
D
Eigenvalues
Variance Explained
Cumulative Variance Explained
PC
E
K-means
WSS
Difference in Slope
No. of Groups
F
Insensitive
Sensitive
Chromophobe-like

FIGURE S10

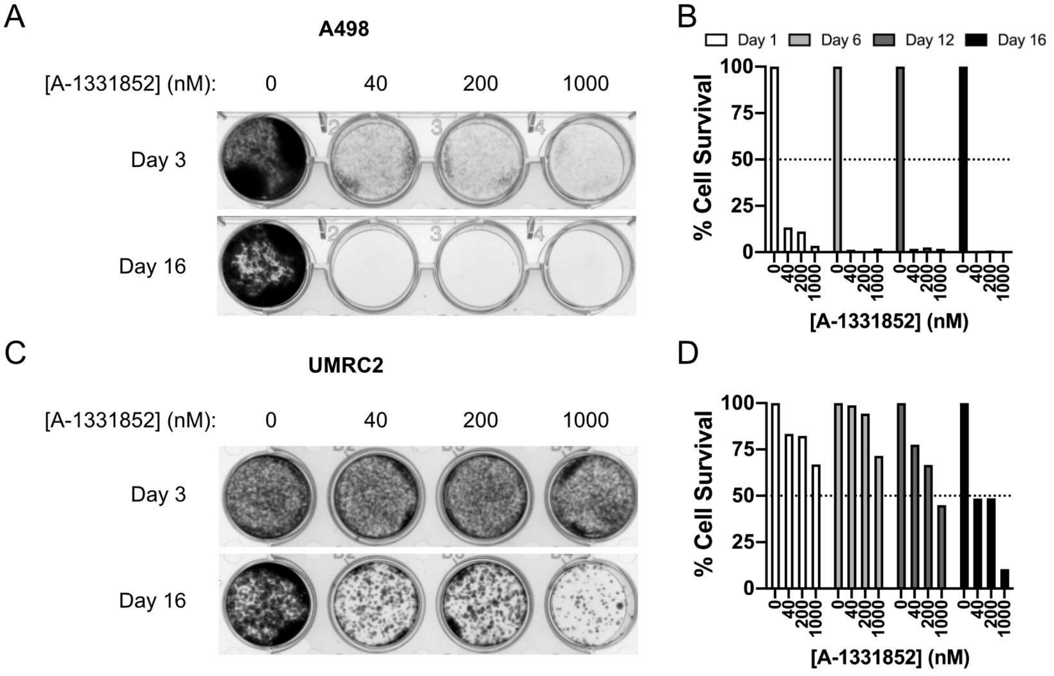

A
A498
[A-1331852] (nM): 0 40 200 1000
Day 3
Day 16
B
Day 1
Day 6
Day 12
Day 16
% Cell Survival
100
75
50
25
0
0 40 200 1000
[A-1331852] (nM)
C
UMRC2
[A-1331852] (nM): 0 40 200 1000
Day 3
Day 16
D
% Cell Survival
100
75
50
25
0
0 40 200 1000
[A-1331852] (nM)

FIGURE S11

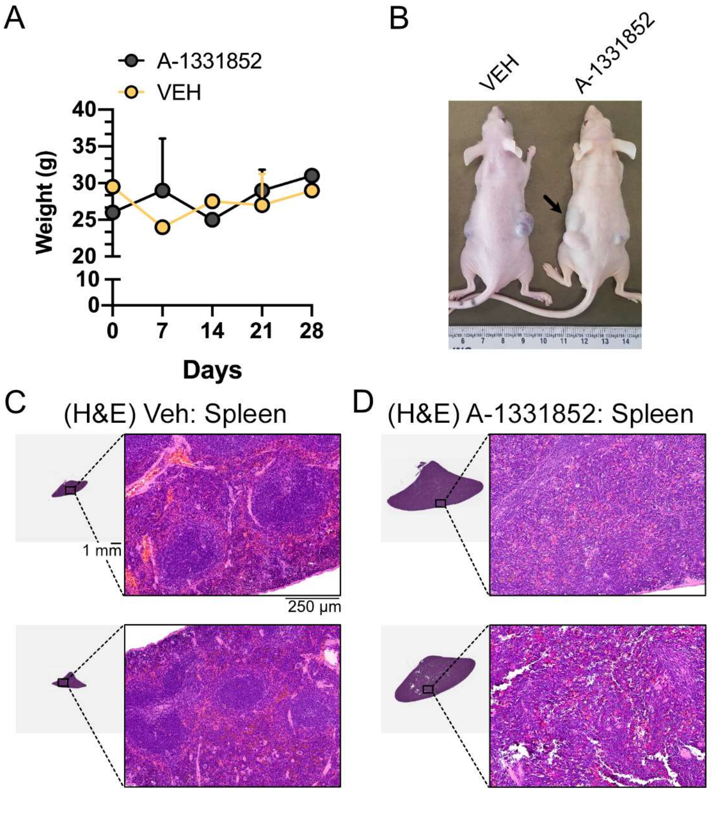
A
A-1331852
VEH
Weight (g)
40
35
30
25
20
10
0
0
7
14
21
28
Days
B
VEH
A-1331852
C
(H&E) Veh: Spleen
1 mm
250 µm
D
(H&E) A-1331852: Spleen

FIGURE S12

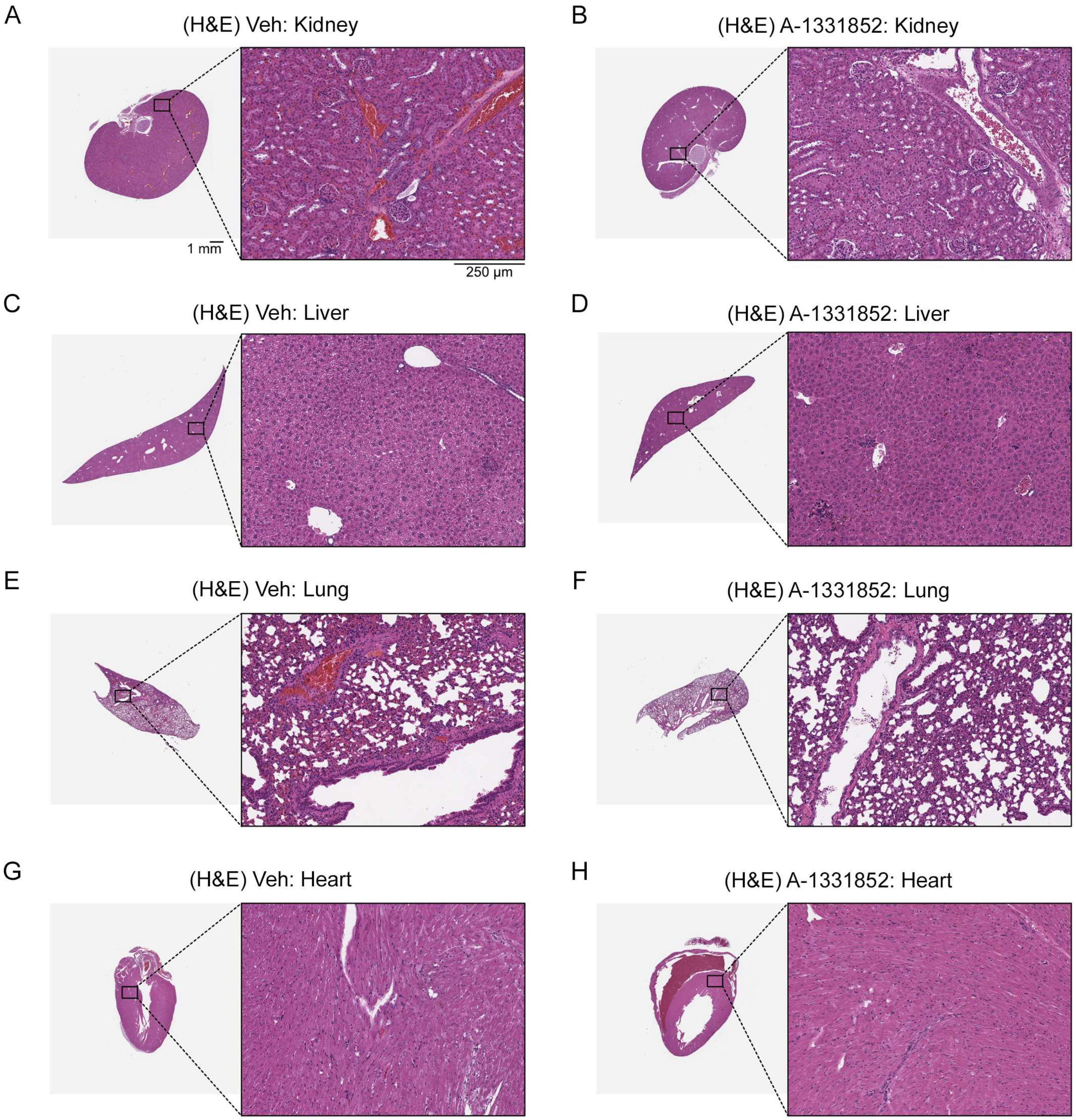
A
(H&E) Veh: Kidney
1 mm
250 µm
B
(H&E) A-1331852: Kidney
C
(H&E) Veh: Liver
D
(H&E) A-1331852: Liver
E
(H&E) Veh: Lung
F
(H&E) A-1331852: Lung
G
(H&E) Veh: Heart
H
(H&E) A-1331852: Heart

FIGURE S13

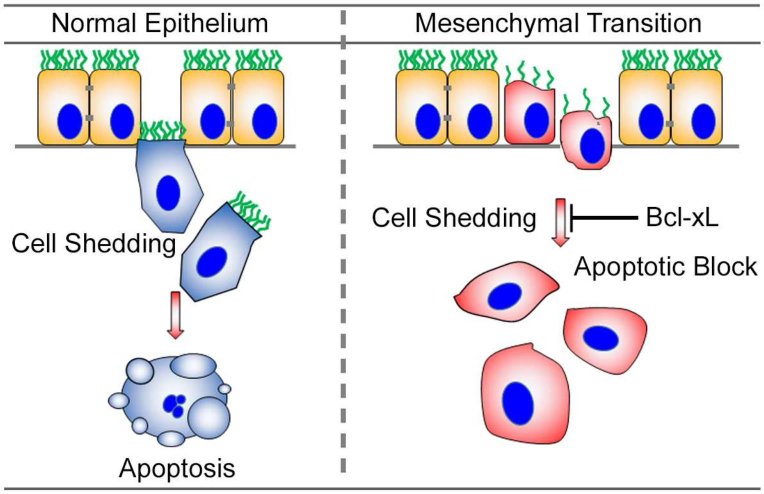
Normal Epithelium
Mesenchymal Transition
Cell Shedding
Apoptosis
Cell Shedding
Bcl-xL
Apoptotic Block
