## Supplemental Figure Legends for "A Mesenchymal Tumor Cell State Confers Increased Dependency on the BCL-X_L_ Anti-apoptotic Protein in Kidney Cancer"

### SUPPLEMENTARY FIGURE LEGENDS

***fig. S1. Pharmacological Characterization of ABT-263 in ccRCC cells.*** (**A** and **B**) Percent cell death, as determined using XTT assays (**A**) and $IC_{50}$ values modeled using standard regression of log(dose) versus response (**B**), in the indicated ccRCC cell lines. In (**A**) data represents mean±S.D. n ≥ 3. (*) represents $IC_{50}$ were calculated by extrapolation because cytotoxicity was < 50% even at the highest treated concentration.

***fig. S2. Pharmacological Characterization of the BCL-X$_L$ inhibitor (A-1331852) and the BCL-2 inhibitor (ABT-199) in ccRCC cells.*** (**A** and **B**) $IC_{50}$ values modeled using standard regression of log(dose) versus response, from at least 3 independent measurements in the indicated ccRCC cell lines, either using the BCL-X$_L$ inhibitor A1331852 (**A**) or the BCL-2 inhibitor ABT-199 (**B**). ND represents Not Determined, because the cytotoxicity was below 50% death, relative to untreated control.

***fig. S3. Measurement of Apoptotic Cell Death upon Pharmacological BCL-X$_L$ Inhibition.*** (**A** to **D**) Flow cytometric analysis to compare AnnexinV-FITC (AnnV) versus Propidium Iodide (PI) staining in CAKI-2 (**A**), A-498 (**B**), UMRC-6 (**C**), and UMRC-2 (**D**) cells that were treated with A-1331852 or DMSO (Untreated control), as indicated. In (**A**) cells were treated with 10 nM A-1331852 for 16 hours; whereas, in (**B**), (**C**), and (**D**) cells were treated with 100 nM A-1331852 for 36 hours.

***fig. S4. Drug-drug Interaction Analysis of A-1331852 and Doxorubicin in ccRCC lines.*** (**A** and **B**). Drug-drug interactions, as measured using the SynergyFinder application package (**A**) and percent cell viability, as determined by XTT (**B**), in UMRC2 cells that were treated with the indicated concentrations of the BCL-$X_L$ inhibitor A-1331852 and Doxorubicin for 72 hours. (**C** and **D**) Drug-drug interactions, as measured using the SynergyFinder application package (**C**) and percent cell viability, as determined by XTT (**D**), in OSRCC2 cells that were treated with the indicated concentrations of the BCL-$X_L$ inhibitor A-1331852 and Doxorubicin for 72 hours. In (**B**) and (**D**), data represents mean±S.D, n = 3.

***Fig. S5. Contribution of BCL-$X_L$ and BCL-2 Expression in A-1331852 Response.*** (**A**) Pearson correlation coefficient of BCL-$X_L$ dependency, as indicated by the DEMETER score, and *BCL2L1* mRNA expression, as measured by RNA-Seq. Cells, which were used in the Achilles dependency analysis (described in Fig. 1) were annotated by lineage. (**B** to **E**) Immunoblot analysis (**B**), densitometric quantification using ImageJ (**C** and **D**), and statistical analysis of the ratio of BCL-$X_L$:BCL-2 abundance (**E**), in the indicated ccRCC cells, which were annotated as “Sensitive”, “Intermediate”, or “Insensitive” based on their response to acute BCL-$X_L$ inhibition (described in Fig. 2 and fig. S2). In (**E**), ns = not significant.

***fig. S6. Contribution of the p53 Pathway in A-1331852 Response.*** (**A**) Annotated genotypes of the indicated cell lines, as described in the Broad Institute’s Cancer Cell Line Encyclopedia. (**B**) Immunoblot analysis in the indicated ccRCC cells, which were annotated as “Sensitive”, “Intermediate”, or “Insensitive” based on their response to acute

BCL-$X_L$ inhibition (described in Fig. 2 and fig. S2), after treatment with Doxorubicin for 6 hours.

***fig. S7. Role of Mesenchymal State in Driving BCL-$X_L$ versus BCL-2 Dependence.*** (**A**) CD44 levels, as determined by flow cytometry, in the indicated BCL-$X_L$ inhibitor Sensitive and Insensitive cell lines. (**B** and **C**) Cell viability, relative to untreated DMSO controls, in UMRC-2 cells that were treated with 10 ng/ml TGFβ for 3 days, and then cultured in the presence of the indicated concentrations of either A-1331852 (**B**) or ABT-199 (**C**) for 7 days. Cell viability, as measured by CellTiter-Glo, was normalized for batch effects and compared using linear regression ($n \geq 3$, dotted line represents best fit, p-values indicated are for difference in slopes)

***fig. S8. Impact of pVHL Status on BCL-$X_L$ Dependence in ccRCC cells.*** (**A** to **D**) Immunoblots (**A**), CD44 expression, as measured by flow cytometry (**B**), and percent cell death, relative to DMSO-treated control cells, in cell lines that were treated with ABT-263 (**C**) or A-1331852 (**D**), as indicated, for 3 days. Cell proliferation was determined using XTT in the indicated ccRCC lines that were lentivirally transduced to express pVHL (VHL) or empty vector (Vector). In (**C**) and (**D**), data represents mean±S.D, n = 3.

***fig. S9. Principal Component Analysis Predicts a BCL-$X_L$ Dependent Signature in Human ccRCC Tumors.*** (**A** to **C**) Eigenvalues (**A**), K-means clustering (**B**), and Principal component analysis (**C**) of the entire kidney cancer gene expression dataset mined from TCGA, showing segregation into 3 principal clusters, driven by disease subtype. (**D** to **F**)

Eigenvalues (**D**), K-means clustering (**E**), and Principal component analysis (**F**) of gene expression data from clear cell Renal Cell Carcinomas (KIRC in TCGA) overlaid with the differential gene signature (described in Fig. 4) to identify 3 principal clusters within ccRCC. As shown in (**F**), two of these clusters represent ccRCCs that resemble cellular signatures of Bcl-xL inhibitor sensitive (red) and insensitive (blue) cells. The third cluster represents (likely misannotated) KIRCs that resemble chromophobe tumors in gene expression patterns (dark grey in **F**).

***fig. S10. Kinetics of Cytotoxicity in Response to BCL-X$_L$ Inhibition in ccRCC cells.*** (**A** and **B**) Crystal violet staining (**A**) and Cell viability, as determined by cell counts (**B**), of A-498 following culture in the presence of the indicated concentration of A-1331852 for the indicated times. (**C** and **D**) Crystal violet staining (**C**) and Cell viability, as determined by cell counts (**D**), of UMRC-2 cells following culture in the presence of the indicated concentration of A-1331852 for the indicated times. In (**B**) and (**D**), cells were counted every using the automated ViCell counter (Beckman) and replated in the presence of fresh drug every 3 days.

***fig. S11. Characterization of splenomegaly in A-1331852-treated animals.*** (**A**) Mean body weight of enrolled mice over the duration of A-1331852 treatment (25 mg/kg, twice a day, Oral gavage). (**B**) Photographs of representative tumor-bearing mice following 4-weeks of twice daily administration of 25 mg/kg A-1331852 or sham-vehicle control. Black arrow indicates the enlarged spleen in the A-1331852 treated mouse. (**C** and **D**) H&E

stains of spleens recovered from either sham-vehicle control (**C**) or A-1331852 (**D**) treated mice, as indicated.

***fig. S12. Histological characterization of the impact of A-1331852 treatment.*** Histological analysis of H&E stained sections of kidney (**A** and **B**), liver (**C** and **D**), lung (**E** and **F**), and heart (**G** and **H**) that were recovered from mice following 4-weeks of twice daily administration of 25 mg/kg A-1331852 or sham-vehicle control, as indicated.

***fig. S13. Proposed Model Linking Cell State to BCL-X$_L$ Dependency.*** Schema representing the protective apoptotic mechanisms that lead to cell death in normal cells that are shed from an organized epithelium in a process called "anoikis" (left). Upon transformation, mesenchymal tumor cells select for mechanisms that can counteract cell death pathways. In kidney cancer cells, this counteracting mechanism is dependent on the activity of the BCL-X$_L$ anti-apoptotic protein.
